# *ace2* expression is higher in intestines and liver while being tightly regulated in development and disease in zebrafish

**DOI:** 10.1101/2020.12.24.424209

**Authors:** Ayse Gokce Keskus, Melike Tombaz, Burcin I. Arici, Fatma B. Dincaslan, Afshan Nabi, Huma Shehwana, Ozlen Konu

**Author notes:** Corresponding Author: Ozlen Konu.

## Abstract

Human Angiotensin I Converting Enzyme 2 (*ACE2*) that acts as a receptor for SARS-CoV-2 entry is highly expressed in human type II pneumocytes and enterocytes and similarly in other mammals and zebrafish (*Danio rerio*). The zebrafish genome has a highly conserved, one-to-one ortholog of *ACE2*, i.e., *ace2*, whose expression profile however has not yet been studied during development or in pathologies relevant to COVID-19. Herein, we identified significant development-, tissue- and gender-specific modulations in *ace2* expression based on meta-analysis of zebrafish Affymetrix transcriptomics datasets (n_datasets_=107, GPL1319 in GEO database). Co-expression network analysis of *ace2* revealed distinct positively correlated (carboxypeptidase activity and fibrin clot formation), and negatively correlated (cilia biogenesis/transport and chromatin modifications) STRING network modules. Using additional transcriptomics datasets, we showed zebrafish embryos before 3 days post fertilization (dpf) exhibited low levels of *ace2* that increased significantly until 4 dpf implicating a role for *ace2* in organogenesis. Re-analysis of RNA-seq datasets from zebrafish adult tissues demonstrated *ace2* was expressed highly in intestines, variably in liver, and at lower levels in other organs. In addition, zebrafish females and males showed significant dimorphism in their age-dependent expression of *ace2*, and between ovary and testis where the latter had higher levels. Moreover, we demonstrated *ace2* expression was significantly modulated under different physiological and pathological conditions associated with development, diet, infection, and inflammation. Our findings implicate a novel translational role for zebrafish *ace2* in differentiation and pathologies predominantly found in intestines and liver, in which the effects of SARS-CoV-2 could be detrimental.

## INTRODUCTION

ACE2 and ACE, involved in angiotensin conversion, are integral elements of renin-angiotensin signaling (RAS) in multiple tissues. Recent studies have shown that RAS is not only present in kidney or adrenal glands but also functional in other tissues with significant roles in multiple pathologies including cancer (Cheng and Liu, 2019; Nehme et al., 2019; Pinter and Jain, 2017; Rasha et al., 2019). The role of ACE2 and ACE in the regulation of hypertension is well known (Hamming et al., 2007) and recently ACE2 has become the focus of intense research due to its function as a receptor of SARS-CoV-2 entry into the cell making it a highly sought after target in the prevention and therapy of COVID-19 (Wan et al., 2020). Several mammalian models and recently zebrafish have been used to study *ACE2* expression and/or function.

Recent findings from single-cell RNA-seq studies show that *ACE2* is highly expressed in type II pneumocytes and intestinal enterocytes in mammals, in which SARS-CoV-2 can replicate and disperse within the tissue of origin and possibly systemically (Ziegler et al., 2020). Moreover, gender might be one of the factors leading to variability in *ACE2* activity due to sex-specific differences observed in the severity of viral infection and mortality rates (Jin et al., 2020). Since *ACE2* has roles in multiple tissues and functions in a wide range of physiological and pathological conditions, understanding mechanisms of SARS-CoV-2 infection has proven to be challenging.

Zebrafish *ace2* is found highly conserved in sequence and structure, and exists as the only copy in zebrafish with a duplicated genome (Chou et al., 2006). Furthermore, a recent study phylogenetically comparing genes co-expressed with *ace2* based on a zebrafish single-cell RNA-seq tissue data has revealed RAS signaling is also highly conserved between humans and zebrafish (Postlethwait et al., 2020). However, there is no study yet in the literature investigating the changes in *ace2* expression over zebrafish embryonic and larval developmental stages, and in different whole adult tissues as well as across different genotypes and/or treatments. In particular, potential sexual dimorphism between males and females over age as well as pathologies relevant to SARS-CoV-2 infection warrant further study.

Herein, we have used bioinformatics approaches and the wealth of transcriptomics data in literature to study the expression patterns of *ace2* across various conditions. We first statistically analyzed (two-group) a large collection of microarray datasets from the Affymetrix platform (GPL1319) and identified development-, tissue-, and gender-specific differences in *ace2* expression. Using additional datasets, we then showed that *ace2* levels increased between 3 and 4 days post fertilization (dpf) and then remained high for the remainder of juvenile and adult life of zebrafish although significant differences were observed between males and females during young adulthood. *ace2* exhibited high tissue specificity where it was expressed the most in intestines followed by liver although highly variably. STRING network enrichment analyses prioritized positively and negatively ranked local functional pathways and provided support as well as novel leads for future studies. In addition, we showed that *ace2* was modulated in different models of liver and intestinal pathologies making the zebrafish model appropriate for SARS-CoV-2 associated research, considering the gut being one of the transmission routes and liver a potential end organ of COVID-19 pathogenesis (Lin et al., 2020).

## METHODS

### Collection and processing of Affymetrix Zebrafish Chip datasets (GPL1319)

Raw .CEL files for all available datasets for GPL1319 were obtained from NCBI GEO database (Barrett et al., 2013). Groups containing at least two samples were used for comparisons; and datasets with no comparable groups were excluded (n_datasets_ = 107). Normalization of .CEL files for each study was performed, separately, using rma in *affy* package (Gautier et al., 2004). From 107 datasets, 344 two-group comparisons were manually generated with following rules: 1. each experimental group was compared to its corresponding control group; 2. experimental groups with double mutation or morpholino treatment were compared to each of the single mutation/morphant group; 3. only control (or wild type) groups from different tissues, developmental time points or gender were compared with each other; 4. for time series, all other groups were compared to the group with the earliest time point only. Differential expression analysis using limma was conducted for each of the manually generated two-group comparisons with the normalized datasets (Ritchie et al., 2015). FDR and logarithmically (log2) transformed fold change (logFC) thresholds were set stringently (adj. p value < 0.05 and abs(logFC) > 0.5).

### Identification of tissue-specific expression patterns of *ace2* in zebrafish adult tissues

Raw data for selected tissue datasets (Table S1) were obtained from SRA (Chen et al., 2020a) and analyzed using Seven Bridges Cancer Genomics Cloud (CGC; https://www.cancergenomicscloud.org/). Reads were aligned using Star Alignment tool (Dobin et al., 2013); and HTSeq tool (Anders et al., 2015) was used to retrieve count data. Raw count data were normalized using RPKM function from *edgeR* package in R (McCarthy et al., 2012). Tau was used as the tissue specificity index as described previously (Kryuchkova-Mostacci and Robinson-Rechavi, 2017).

### Dataset collection for sexual dimorphism, and intestinal and liver development and disease

Zebrafish transcriptomics datasets selected from NCBI GEO or Expression Atlas (Papatheodorou et al., 2020) according to their relevance to sexual dimorphism, liver and gut development and disease/treatments were summarized in Table S2. Series matrix files for datasets of GPL14664 (GSE113241 (Alvarez-Rodriguez et al., 2018), GSE112272 (Jia et al., 2019), GSE73233 (Forn-Cuni et al., 2017), and GSE100583 (Holden and Brown, 2018)) and RPKM normalized data of GSE74244 (Aramillo Irizar et al., 2018), GSE118076 (San et al., 2018), and GSE24616 (Domazet-Loso and Tautz, 2010) were obtained from GEO and used for statistical analysis after logarithmic (base 2) transformation. The expression of *ace2* in the E-ERAD-475 dataset was obtained from Expression Atlas (White et al., 2017). ANOVA followed by multiple test correction (Tukey’s HSD) were used for statistical analyses as indicated in the Fig. legends. Raw count data of GSE82200 (Koch et al., 2018), GSE83195 (Schall et al., 2017) and GSE123439 (Chen et al., 2020b) were obtained from NCBI GEO; *Deseq2* package was used for the differential gene expression analysis; and Regularized log (Rlog) normalized expression values were used in visualization (Love et al., 2014).

### Enrichment Analysis

A Pearson’s correlation coefficient of *ace2* (r_*ace2*_) was calculated between logFC values of *ace2* and those of every other gene across all comparisons performed with GPL1319 datasets. r_*ace2*_ also was calculated for each gene based on log2 transformed expression values for larval development, using GSE24616 (2 dpf-8 dpf) and GSE38575 (EtOH treated samples only; 2dpf - 7dpf) datasets; for intestine, using GSE35889 and GSE12189 (GFP+ samples only); and for liver, using GSE74244 (liver samples only) and GSE100583, separately (Table S2). r_*ace2*_ value with minimum p value was selected for multiple probe sets with the same Ensembl ID. Ensembl IDs along with r_*ace2*_ values were used for identifying local network clusters using STRING enrichment analysis with values for each of the above datasets (Szklarczyk et al., 2015). Comparative Network Enrichment Analysis (CNEA) of Sting local networks were performed separately for development, intestine and liver datasets. Enriched local networks with the largest gene set was selected in the case of multiple local networks present with the same name. Significant networks were clustered and visualized in Cytoscape (Shannon et al., 2003); where each node represented a local network, colored according to STRING enrichment score (the mean enrichment score in the case of liver and intestine comparisons), having edge weights determined by the number of shared genes with other connected networks. Selected local networks were visualized in detail, where each node on the PPI network obtained from STRING local network data referred to a gene and was colored according to r_*ace2*_. In addition, GO term enrichment analysis was performed for liver datasets using *clusterprofiler* package in R (Yu et al., 2012).

## RESULTS

### Analysis of zebrafish GPL1319 datasets reveal conditions in which ace2 expression is significantly modulated

Upon re-analysis of publicly available datasets from the GPL1319 platform (n=107 with 344 pairwise group comparisons), *ace2* expression was found differentially expressed in 19 pairwise group comparisons from 11 different datasets relevant to sexual dimorphism, estrogen signaling, embryogenesis, and liver and gut development (Table 1) (Froehlicher et al., 2009; Heiden et al., 2008; Hu et al., 2013; Jacob et al., 2015; MacInnes et al., 2008; Okuda et al., 2010; Rogers et al., 2011; Small et al., 2009; Soni et al., 2013; Stuckenholz et al., 2009; Thakur et al., 2014). STRING local network clusters obtained using r_*ace2*_ calculated with logFC values from all comparisons (n=344) revealed that genes positively correlated with *ace2* were enriched in carboxypeptidase activity, intestinal hexose absorption, villin and keratin, and fibrin clot formation associated sub-groups (Figs. 1, S1A) whereas those negatively correlated belonged to cilium assembly, microtubule organization pathways, and chromatin modifying enzymes (Figs. 1, S1B). Ten genes were represented in GPL1319 platform out of 28 RAS pathway genes (Postlethwait et al., 2020); and eight of them were found to be positively correlated with *ace2*, with a correlation coefficient ranging from 0.46 to 0.85.

**Table 1:**
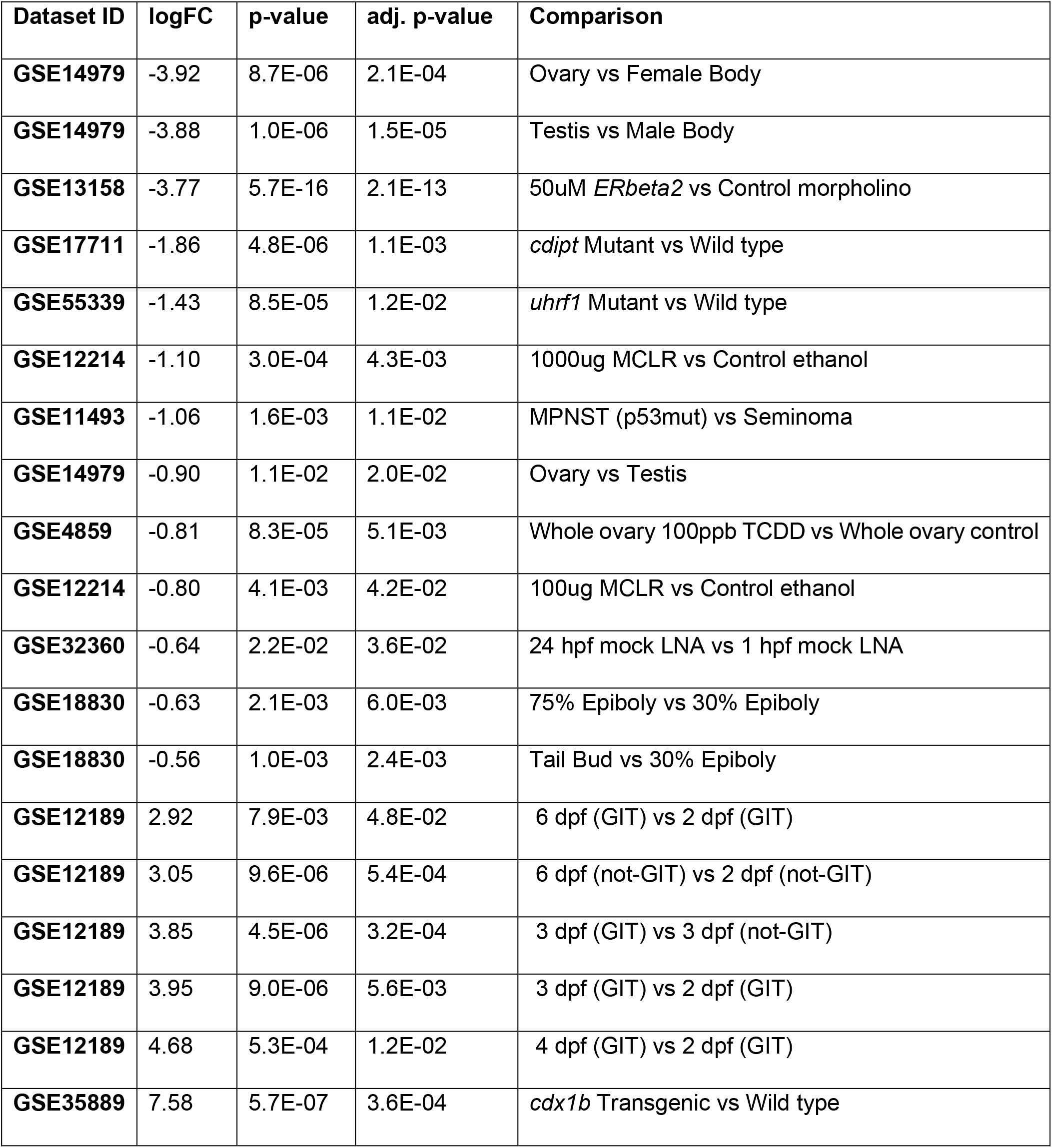
GPL1319 datasets in which zebrafish *ace2* with a probe set ID “Dr.20290.1.A1_at” was significantly differentially expressed (FDR (adj. p-value) < 0.05 and abs(logFC) > 0.5). MCLR, MPNST, TCDD, LNA, GIT, refer to microcystin-LR, malignant peripheral nerve sheath tumor, 2,3,7,8-Tetrachlorodibenzo-p-dioxin, locked nucleic acid, and gastrointestinal tract.

**Fig. 1.**
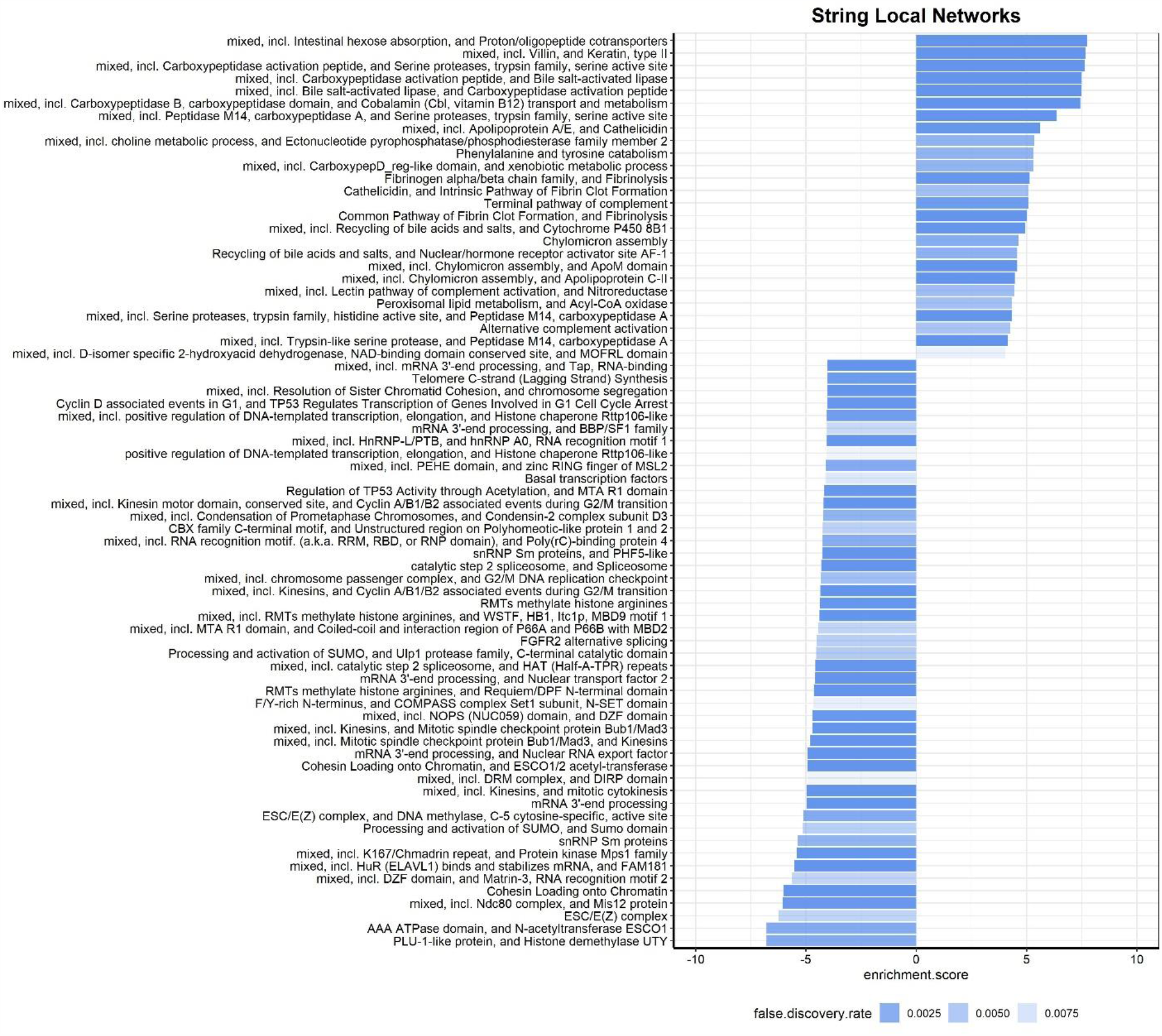
Bar plots of significant STRING local network enrichment scores obtained from analysis of r_*ace2*_ values across GPL1319 datasets (n_dataset_ = 107; n_comparison_ = 344; abs (enrichment score) > 0.4).

### *ace2* expression in zebrafish is developmentally regulated

Analysis of GPL1319 datasets revealed *ace2* expression decreased throughout gastrulation (GSE18830, GSE32360; Table1), which was supported by re-analysis of GSE24616 (Fig. 2A) and E-ERAD-475 (Fig. S1C). In addition, we found *ace2* was lowly expressed in embryogenesis and started increasing at 3 dpf and thereafter until 4 dpf, after which was steadily expressed at high levels (Figs. 2A, S1C). Analysis of yet another dataset (GSE38575) supported this finding of an increase in *ace2* expression after 3 dpf (Fig. 2B). STRING enrichment scores based on r_*ace2*_ between *ace2* and every other gene obtained separately for GSE24616 (2dpf - 8dpf) and GSE38575 (untreated; 2dpf - 7dpf) datasets were significantly correlated with each other (r = 0.963, p-value = 7.63e-127; Fig. 2A-C). Network clustering helped us define pathways modulated at the time of *ace2* induction in early zebrafish larvae. Accordingly, networks enriched with genes positively correlated with *ace2* belonged to pathways related to intestinal hexose absorption, Vitamin D metabolism, carboxypeptidases, interleukin signaling, phenylalanine/tyrosine catabolism, peptide ligand binding and dopamine receptors, phototransduction, and common fibrin clot/fibrinolysis (Figs. 2D, S2). On the other hand, networks enriched with negatively correlated genes included terms such as chromosome segregation, DNA double-strand break and replication, homeobox, alternative splicing, and endothelial cell proliferation (Figs. 2D, S3, Table S3).

**Fig. 2.**
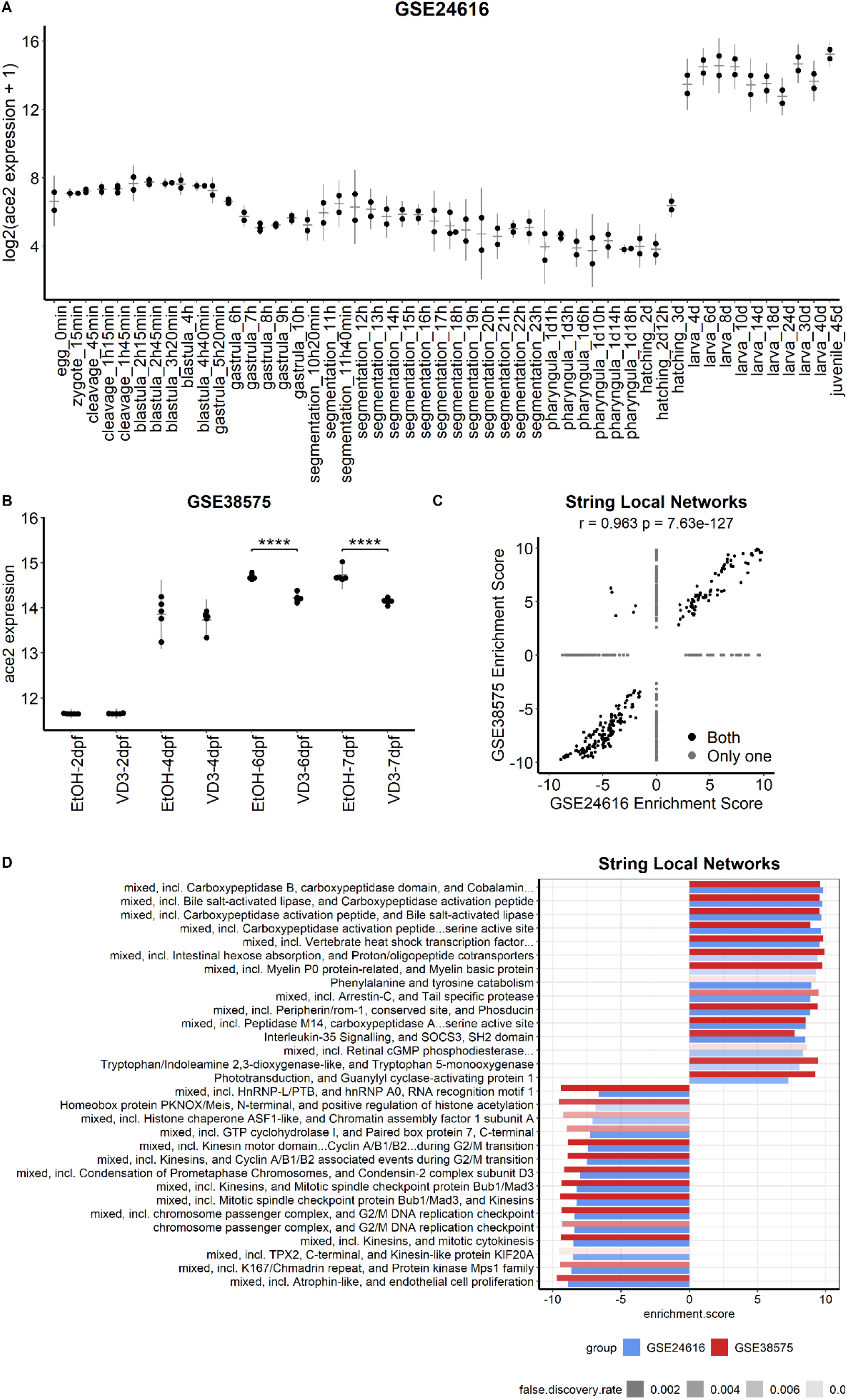
*ace2* expression patterns during developmental stages from GSE24616 (A) and GSE38575 (B); and correlation between significantly modulated STRING local networks with *ace2* expression during development (C). Top 15 positively and negatively enriched networks with *ace2* expression during development (D).

### *ace2* expression in zebrafish is sexually dimorphic

One of the most prominent findings of the GPL1319 dataset meta-analysis was the identification of sexual dimorphism in zebrafish *ace2* expression as shown in Table 1 (GSE14979; Table 1). Interestingly, gonads had lower *ace2* expression when compared with the rest of the body regardless of sex, while testis had relatively higher *ace2* levels than the ovary (GSE14979; Fig. S4A). Moreover, *ace2* expression decreased in the presence of a morpholino for *ERBeta2* (*esr2a*) (GSE13158; Table 1), previously shown to be essential for female sexual maturation and early follicle generation (Lu et al., 2017; Wu et al., 2020). We then re-analyzed the GSE24616 dataset for *ace2* expression in adult zebrafish through time and found *ace2* was highly expressed in juveniles and early adulthood (90 dpf) (Fig. 3A). Moreover, males exhibited high *ace2* expression regardless of age while females showed a steady decrease over aging until 9 months (Fig. 3A; adj. p-value <0.0001 (age), adj. p-value <0.0001 (gender), adj. p-value = 0.43 (interaction)); however, after 9 months ace2 levels were indistinguishable between males and females. Results obtained from another dataset, namely, GSE123439, was consistent with the observed significance of sexual dimorphism in gonads (Fig. S4B).

**Fig. 3.**
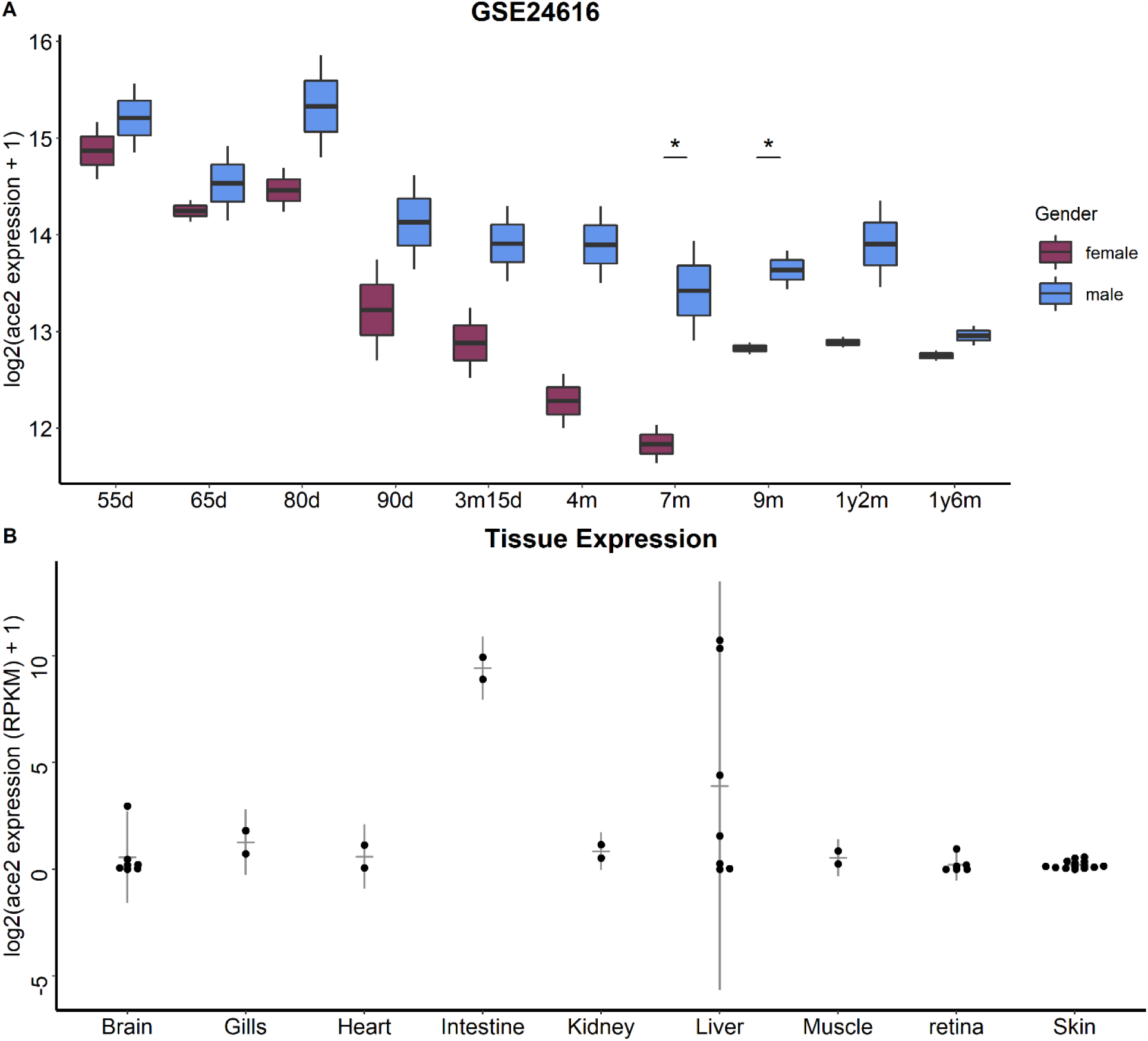
Sexual dimorphic expression pattern in adulthood based on GSE24616 (Simple effect analysis following Two-Way ANOVA results were represented. *, p-value < 0.05) (A). Tissue specific expression *of ace2* (log2(RPKM+1)) in intestines (731.1) and liver (432.8) and the relatively low expression in gills (1.57), brain (1.08), kidney (0.83), heart (0.62), and muscle (0.49) in the six month old zebrafish cohort (B).

### *ace2* exhibits the highest expression in intestines

In our meta-analysis of GPL1319 datasets, *ace2* expression also increased the most in intestines of the zebrafish overexpression model of *cdx1b*, an intestinal differentiation related transcription factor, under squamous epithelial cell specific krt5 promoter (GSE35889, Table 1). *ace2* expression increased in GFP+ gastrointestinal tract (GIT) cells of transgenic (Tg(XlEef1a1:GFP)s854) zebrafish larvae at and after 3 dpf compared to 2dpf (GSE12189, Table1; Fig. S5A) while in GFP- (non-GIT) cells, *ace2* was significantly upregulated at 6 dpf only. Since *ace2* expression also increased in GFP+ cells when compared to GFP-cells at 3 dpf (GSE12189, Table1; Fig. S5A), we further performed comparative network enrichment analysis (CNEA) of String local networks for GSE12189 and GSE35889. Network enrichment scores of the abovementioned datasets were highly correlated (r = 0.798, p-value = 6.49e-12; Fig. S5B); and top common networks included those of intestinal hexose absorption, chylomicron assembly, i.e., a key mechanism for lipid transport in the intestines, common pathway of fibrin clot formation and fibrinolysis, fatty acid degradation and carboxypeptidases (Figs. S5C, S6). On the other hand, negatively correlated genes were enriched in networks of striated muscle contraction and mRNA splicing (Figs. S5C, S6).

To support the role of ace2 in intestinal function and differentiation we next investigated the tissue specificity of *ace2* expression across multiple datasets we complied from RNA-seq experiments (Fig. 3B; Table S1). Intestine and liver exhibited relatively high *ace2* expression in comparison with brain, gills, and kidney in that descending order (Fig. 3B). Accordingly, zebrafish *ace2* expression was found to have a tissue specificity index value (Tau) of 0.99. Moreover, increases in *ace2* expression were steady over time in late zebrafish larvae with functional intestines (Fig. S7A).

### *ace2* expression is altered with inflammation

From an intestinal disease perspective, *ace2* was found to be downregulated in the zebrafish model of Short Bowel Syndrome (SBS) (Fig. S7B) which may implicate a possible link between *ace2* expression and inflammatory response (Johnson et al., 2018; Mutanen et al., 2019; Schall et al., 2017). We, therefore, re-analyzed selected expression datasets associated with different inflammatory stimuli in zebrafish. No significant alteration in *ace2* expression was observed in response to bacterial colonization (Conventionalization (CONVD), *Exiguobacterium* (Exi) and *Chryseobacterium* (Chrys)), or Spring Viraemia Carp Virus (SVCV) infection alone or in the presence of immunostimulants (GSE113241, Fig. 4A; GSE73223, Fig. S7C). However, *ace2* expression significantly increased with immunosuppression in the zebrafish *myd88* knock-out model showing significant effects for genotype and treatment with a significant interaction in between these factors (adj. p-value = 4e-07 (Genotype), adj. p-value = 0.03 (Germ), adj. p-value = 0.03 (interaction); Fig. 4A). On the other hand, *ace2* expression decreased with SVCV infection after pre-treatment with β-glucan but not with exposure to lipopolysaccharides (LPS) or Polyinosinic:polycytidylic acid (PolyI:C) exposure (adj. p-value = 0.01 (SVCV), adj. p-value = 0.06 (pre-treatment), adj. p-value = 0.04 (interaction); Fig. 4B). LPS treatment was not effective in liver, muscle, or kidney, either (Fig. S7C).

**Fig. 4.**
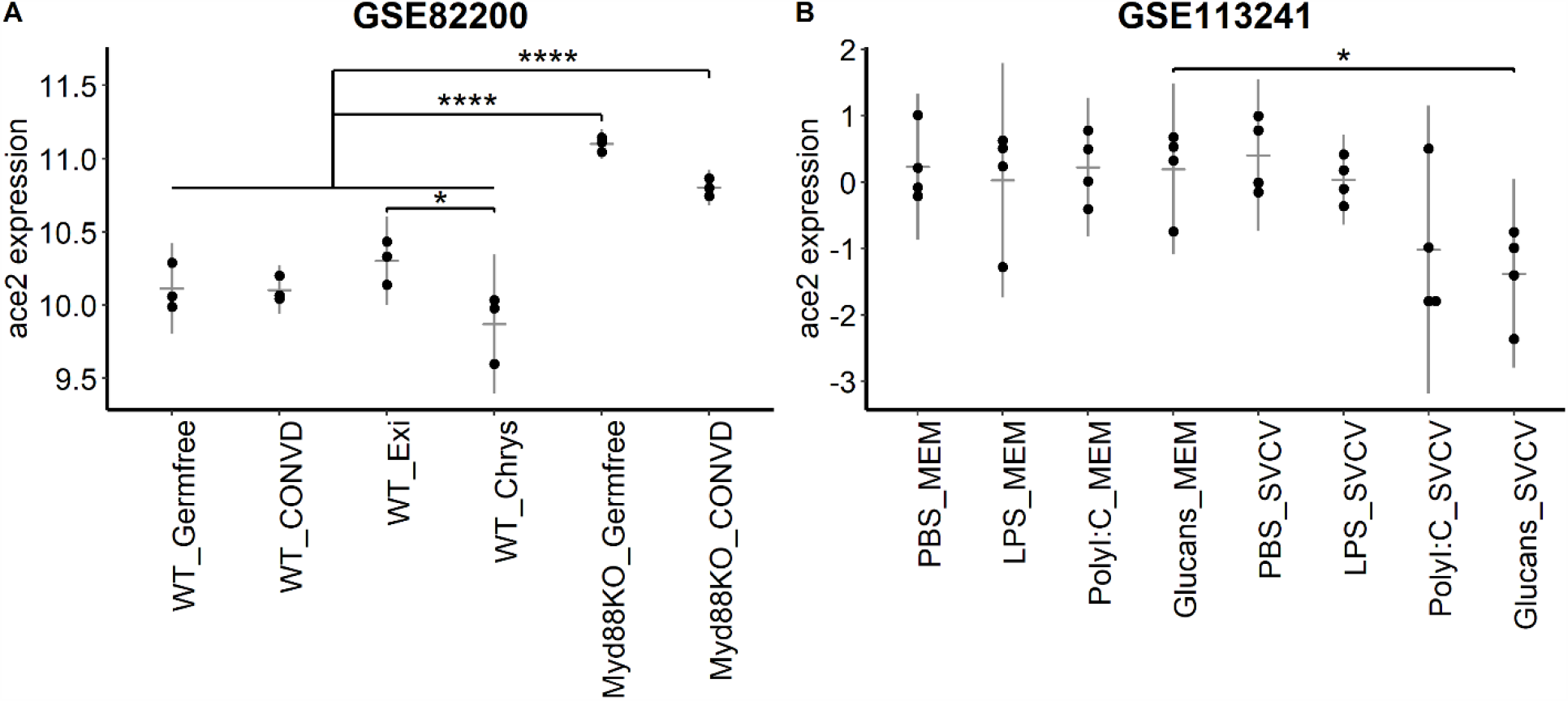
*ace2* expression during immune response in *myd88* knock-out (Myd88KO) model with bacterial colonization (Conventionalization (CONVD), *Exiguobacterium* (Exi) and *Chryseobacterium* (Chrys)) (Tukey HSD corrected One-way ANOVA results were represented. *, adj. p-value < 0.05; ****, adj. p-value < 0.0001) (A); in viral infection (SVCV: Spring Viraemia Carp Virus) after pre-treatment with β-glucans, lipopolysaccharide (LPS), or Polyinosinic:polycytidylic acid (PolyI:C) (Simple effect analysis following Two-Way ANOVA results were represented. *; p-value < 0.05) (B).

### *ace2* expression is highly variable in liver and associated with diet and liver disease

Since *ace2* exhibited the second highest yet bimodal expression levels in liver (Fig. 3B; Fig. S8A), we performed GO annotation of genes with high and low r_*ace2*_ values in GSE74244 (the largest RNA-seq zebrafish cohort for zebrafish adult liver tissue in GEO database). Significant GO biological processes of genes whose expression levels were positively correlated with that of *ace2* were enriched in metabolism, and in particular, that of the lipid (Fig. S8B) whereas those negatively correlated included immune response (Fig. S8C). We showed that liver expression was not sexually dimorphic and could vary regardless of strain and gender using another dataset (GSE100583, Fig. S9A). Results of GO enrichment analysis of GSE100583 were similar in terms of positively correlated genes yet cilium organization and establishment of cell polarity were found among significantly negatively correlated pathways (Fig. S9B, C). CNEA of STRING Local Networks between these two independent datasets were highly concordant for a subset of pathways (Fig. 5A, Fig. S10A) and included networks of genes with positive r_*ace2*_ such as intestinal brush border proteins (villin and keratin) as well as interferon stimulated genes (ISG15 antiviral mechanism), carboxypeptidases, and respiratory electron transport and oxidative phosphorylation (Figs. 5A, S10B). Erythropoietin and hemoglobin network was among those enriched with negative r_*ace2*_ values in both datasets (Figs. 5A, S10B). However, apolipoprotein A/E genes, known to be functional in fat metabolism and immune response, although enriched in both datasets acted in the opposite direction (Fig. 5A).

**Fig. 5.**
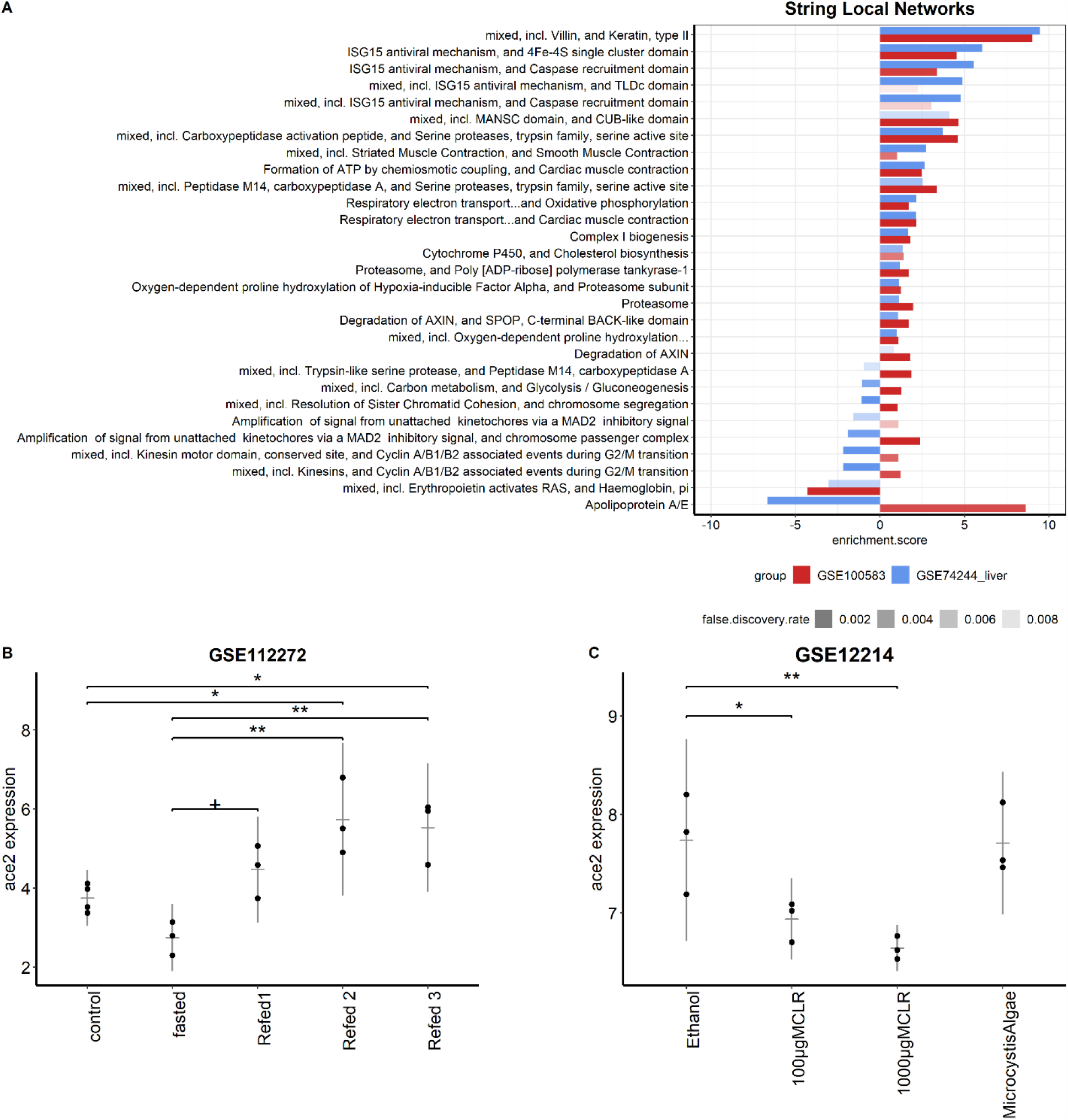
Bar plots of significant STRING local network enrichment scores for GSE74244 and GSE100583 liver datasets (A); *ace2* expression in zebrafish livers during fasting and refed conditions (B); in microcystin-LR (MCLR) exposure (*, p-value < 0.05; **, p-value < 0.01) (C).

We further investigated *ace2* expression in zebrafish under conditions of fasting and refeeding in the liver and found *ace2* expression increased when refed after 3 weeks of fasting (Fig. 5B) yet it was not altered when given a high carbohydrate diet (GSE8856 (Robison et al., 2008); logFC = 0.24, adj. p-value = 0.51 (females) and logFC = -0.67, adj. p-value = 0.99 (males)) or under the condition of overfeeding (GSE48806 (unpublished data); logFC =-1.76, adj. p value = 0.26). In further support of involvement of liver, *ace2* expression decreased in the zebrafish non-alcoholic fatty liver disease (NAFLD) model (GSE17711; Table 1) and in response to microcystin-LR, previously shown to cause liver damage (GSE11214; Table1, Fig. 5C) (Rogers et al., 2011).

## Discussion

Zebrafish as a model organism is widely used to understand disease mechanisms due to its high sequence/functional similarity with humans as well as the availability of a wide range of zebrafish genetic models (Gomes and Mostowy, 2020; Logan et al., 2018; Mickiewicz et al., 2019; Shehwana and Konu, 2019; Willis et al., 2016). ACE2, identified as the receptor allowing entry of SARS-CoV-2 to the host cell, has a single ortholog in zebrafish with 65% identity (Chou et al., 2006); and the RAS pathway in zebrafish has been shown to be similarly active as in humans (Postlethwait et al., 2020). However how zebrafish *ace2* expression changes in embryonic/larval development or disease remains unexplored; hence, we investigated this using meta-analysis and/or functional network enrichment approaches with respect to various contexts including age, gender, tissue specificity and pathologies with relevance to COVID-19.

We have demonstrated that *ace2* is expressed at low levels during early stages of larval development yet starts increasing in expression during liver and intestinal differentiation (Kimmel et al., 1995). Developmental studies in mice report *Ace2* expression at E12.5 day with an increasing trend over time in kidneys, lung, brain and heart (Song et al., 2012). *ACE2* is also expressed in human intestines, gonads, kidneys, heart, adipose tissue, lung, and liver in accord with its multiorgan functionality (Musavi et al., 2020; Wang et al., 2020). Herein based on re-analysis of multiple transcriptomics datasets (both embryonic and adult), *ace2* has been shown to be tissue-specifically expressed in zebrafish adult intestines as in humans (Musavi et al., 2020). The potential role of ace2 in zebrafish intestinal development is further supported by a drastic increase in *ace2* expression (logFC = 7.58) in the *cdx1b* transgenic exhibiting reduced squamous epithelium at the expense of increased intestinal differentiation. This finding supports that *cdx1b* could be a transcriptional regulator of *ace2* in zebrafish as *Cdx2* is in mice (Chen et al., 2020a). Indeed, *cdx1b* is the functional ortholog of human *CDX2*, an important transcription factor with known functions during digestive system development in humans (Cheng et al., 2008; Flores et al., 2008; Silberg et al., 2000). In addition, fish larvae with a mutation in *uhrf1* gene, reported to cause non-functional intestines, also exhibit lower levels of *ace2* further supporting a potential role for *ace2* in intestinal development (Table 1) (Ganz et al., 2019; Marjoram et al., 2015).

Herein, CNEA of STRING local networks (Yildiz et al., 2013) have proven to be an effective strategy demonstrating that *ace2* expression is a highly co-expressed component of carboxypeptidases; this supports the reports proposing serine protease family of genes as a target for COVID-19 treatment (reviewed in (Vargas et al., 2020; Yao et al., 2020)). Other networks that warrant further attention in the context of SARS-CoV-2 infection include fibrinolysis (positive r_*ace2*_) since COVID-19 severity is associated with dysregulation of clot formation (reviewed in (Coccheri, 2020)). Our findings also suggest that zebrafish can serve as a potential model organism for testing effects of *ace2* over- or under-expression as well as candidate drugs (e.g., peptidase inhibitors) on enzymatic and complement activity in larval and adult intestines under SARS-CoV-2 infection(Donmez et al., 2020).

Interestingly, several studies have previously reported a higher *ACE2* expression in cilia of well differentiated epithelial cells compared to undifferentiated cells (Jia et al., 2005; Lee et al., 2020b; Wan et al., 2020). *ACE2* expression level in cilia is also shown to correlate with SARS-CoV infection (Lee et al., 2020b), and infection with coronavirus causes loss of cilia (Wan et al., 2020). Most recently ACE2 has been shown to be expressed in motile cilia in airway epithelial cells in humans (Lee et al., 2020a). In accord, herein we found genes negatively correlated with *ace2* were enriched in cilia, intraflagellar transport, and microtubule organization, in particular, during gastrointestinal development and intestinal differentiation. Further studies are needed to understand the dynamicity between *ace2* expression and cilium assembly; and zebrafish provides an effective translational model for testing potential drugs affecting intraflagellar transport in the context of SARS-CoV-2 infection (Duan et al., 2020).

Consistently, our analyses of r_*ace2*_ in zebrafish have pointed to the enrichment of respiratory chain related networks. In support of this, *Ace2* knockout mice has been reported to exhibit disrupted mitochondrial function in pancreatic islets and skeletal muscle cells (Cao et al., 2019; Shi et al., 2018). Transcriptomic analyses with human cornea, retina and lung datasets propose *ace2* is highly co-expressed with mitochondrial genes (Yuan et al., 2020). Indeed, Ang II is known to activate NADPH and can stimulate an increase in reactive oxygen species (ROS) in renal cells while *ACE2*, as a negative regulator of Ang II, has a protective role against ROS (Gava et al., 2009; Gwathmey et al., 2010; Kim et al., 2012). *Ace2* knockout mice also exhibit increased insulin resistance which can lead to hepatic stress along with increased oxidative stress in accordance with Mas knockout mice (Cao et al., 2014; Gallagher et al., 2008; Gwathmey et al., 2010; Song et al., 2020). Zebrafish can serve as a model organism to study the potential link between the modulatory role of *ACE2* in mitochondrial function and adverse effects of diabetes on COVID-19 prognosis.

Our analyses have indicated liver as the second highest *ace2* expressing tissue in zebrafish yet with a clear bimodal pattern regardless of age (6 months – 42 months) or gender. The functional distinction between adult livers expressing *ace2* at lower or higher levels has become a question of interest and our comparative functional enrichment methodologies provide potential answers.

GO enrichment analyses of *ace2*’s positively co-expressed genes reveal the involvement of lipid catabolism as a potential discriminator; in support of this, we have shown that *ace2* levels decrease in response to microcystin-LR, a common form of cyanotoxin that is reported to cause liver damage as well as disruption of lipid metabolism (Liu et al., 2014; Woolbright et al., 2017). This finding may have translational implications and warrants further examination since the rates of severity and morbidity in COVID-19 increase with obesity (BMI>25) (Kruglikov et al., 2020). Indeed, our identification of steadily increased *ace2* expression in zebrafish refed liver after fasting aids into the connection between *ace2* expression and diet. For instance, *Ace2* levels also have been shown to be elevated in high-fat fed mouse and rat (Gupte et al., 2008; Shoemaker et al., 2019; Zhang et al., 2014), however not in the initial response to lipid exposure in zebrafish digestive organs (Zeituni et al., 2016). This suggests that further experiments are needed to identify the time-frame diet can alter *ace2* levels in zebrafish. In our re-analysis, we also have found *ace2* levels decrease in the NAFLD zebrafish model, in accord with mouse data in literature (Cao et al., 2016; Yang et al., 2020). Interestingly an increasing number of studies shows the inverse relation between NAFLD and COVID-19 recovery (Duan et al., 2020; Forlano et al., 2020; Huang et al., 2020; Mahamid et al., 2020). Our findings indicate that increased oxidative phosphorylation as well as altered lipid metabolism, also observed in obesity, might be associated with increased *ace2* levels in liver, an end organ in COVID-19 (Jothimani et al., 2020; Pawlotsky, 2020), and might have a role in modulating the disease severity.

In addition to altered lipid metabolism, interferon pathway modulation (ISG15 antiviral mechanism network) has arisen as a discriminator of low and high *ace2* expressing liver in zebrafish. Interestingly, studies with SARS-CoV-2 reported *ACE2* as an interferon stimulated gene (ISG) in humans (Chua et al., 2020; Ziegler et al., 2020). Liver, an organ continually subjected to food antigens and pathogens from intestines, can respond to such activators of immune response readily with its residential macrophages (Bleriot and Ginhoux, 2019); and this may result in highly variable interferon activity in the liver (reviewed in (Robinson et al., 2016)). Future studies focusing on embryonic and tissue-specific *ace2* knockout and overexpression models can help understand the role of adaptive and innate immune system challenges in zebrafish liver expression variability.

Moreover, a prognostic importance for Vitamin D has been shown in COVID-19 such that vitamin D exposure potentially increases *ACE2* levels as well as reducing the risk for cytokine storm (Benskin, 2020; Bleizgys, 2020; Musavi et al., 2020). Herein we have identified a significant decrease in *ace2* levels in response to Vitamin D treatment during early larval and organ development that can serve as an important model to investigate the mechanism behind the association between Vitamin D levels and *ace2* expression.

The observed decrease in *ace2* expression in the zebrafish SBS model is also intriguing and points to a link between decreased levels of *ace2* and inflammation as widely observed in human SBS patients (Schall et al., 2017). Our re-analysis has shown that zebrafish *uhrf1* and *cdipt* mutants with elevated intestinal inflammation both exhibit a significant decrease in *ace2* expression (Marjoram et al., 2015; Thakur et al., 2014). In human inflammatory bowel disease (IBD) *ACE2* levels exhibit a similar decrease in inflamed ileum yet increases in inflamed colon and rectum (Suarez-Farinas et al., 2020; Verstockt et al., 2020). Interestingly, patients with IBD along with other inflammatory diseases might have lower susceptibility to COVID-19 (Suarez-Farinas et al., 2020; Verstockt et al., 2020). Furthermore, a strong association has emerged between pathways related to COVID-19 and those of IBD (Suarez-Farinas et al., 2020). In support of the link between *ace2* levels and inflammatory pathways, our meta-analysis shows that zebrafish embryos deficient in the key innate immunity regulatory factor *myd88* exhibit altered lipid metabolism while expressing *ace2* highly, regardless of the type of microbiome (Koch et al., 2018). Interestingly, in mouse, *MyD88* is shown to be essential in protection against SARS-CoV through activation of NF-κB pathway upon viral infection (Hirano and Murakami, 2020; Sheahan et al., 2008), which needs to be tested in zebrafish immune deficiency models. Moreover, our bioinformatics analyses show that preexposure to immune suppressive and lipid modulatory agent β-glucans may lower *ace2* levels (AbuMweis et al., 2010) supporting that the increased fatty acid synthesis might inhibit immune response (Yaqoob and Calder, 2007), and possibly *ace2* levels in zebrafish.

Although the mortality risk of COVID-19 is relatively higher in males than females, there is no concurrent evidence to link the sexual dimorphic effect of COVID-19 to the disease mechanism and the role *ACE2* expression might play in humans. Indeed, analysis of human tissue *ACE2* expression has indicated no significant difference between males and females in general while testis is shown to have higher expression than the ovary (Musavi et al., 2020). Our findings in the present study points to a sexually dimorphic expression pattern of *ace2* in zebrafish, which can provide a model for studying gender specific *ace2* expression in response to different drugs to be tested for COVID-19 treatment.

Moreover, *esr2a* along with estrogen level play important roles in zebrafish starting from early developmental stages as well as during sexual maturation; and *esr2a* knockout zebrafish broods exhibit a significantly lower female to male ratio (Lu et al., 2017; Wu et al., 2020). This finding implicates *esr2a*, as a potential modulator of *ace2* expression and can also be studied in other animal models. In addition, the sexual dimorphism observed in *ace2* levels is more prominent before one year of age at which the females are at their most fertile stage may indicate a role for estrogen signaling in regulation of *ace2* expression. Our results showing a decrease in *ace2* expression in the zebrafish models of *esr2a* knock-down and TCDD exposure, an environmental toxicant acting as an estrogen inhibitor, might help explain gender-specific protective influence of estrogen against the COVID-19 infection-mediated or inherent upregulation of *ACE2* expression.

In conclusion, COVID-19 is a multiorgan disease affecting lungs, kidney, intestines, and liver primarily and recent studies suggest possible impact on other tissues including retina and brain (Noris et al., 2020; Stein and Young, 2020; Zaim et al., 2020). Tissue and context-dependent expression of *ace2* in zebrafish correlate well with human, primate, and rodent data as exemplified above. Therefore, zebrafish with its high translational potency can serve as an effective model organism for COVID-19 research while our comprehensive bioinformatics and statistical analyses provide novel physiological and pathological implications to follow up.

## References

AbuMweis, S. S., Jew, S. and Ames, N. P. (2010). beta-glucan from barley and its lipid-lowering capacity: a meta-analysis of randomized, controlled trials. Eur J Clin Nutr 64, 1472–80.

Alvarez-Rodriguez, M., Pereiro, P., Reyes-Lopez, F. E., Tort, L., Figueras, A. and Novoa, B. (2018). Analysis of the Long-Lived Responses Induced by Immunostimulants and Their Effects on a Viral Infection in Zebrafish (Danio rerio). Front Immunol 9, 1575.

Anders, S., Pyl, P. T. and Huber, W. (2015). HTSeq--a Python framework to work with high-throughput sequencing data. Bioinformatics 31, 166–9.

Aramillo Irizar, P., Schauble, S., Esser, D., Groth, M., Frahm, C., Priebe, S., Baumgart, M., Hartmann, N., Marthandan, S., Menzel, U. et al. (2018). Transcriptomic alterations during ageing reflect the shift from cancer to degenerative diseases in the elderly. Nat Commun 9, 327.

Barrett, T., Wilhite, S. E., Ledoux, P., Evangelista, C., Kim, I. F., Tomashevsky, M., Marshall, K. A., Phillippy, K. H., Sherman, P. M., Holko, M. et al. (2013). NCBI GEO: archive for functional genomics data sets--update. Nucleic Acids Res 41, D991–5.

Benskin, L. L. (2020). A Basic Review of the Preliminary Evidence That COVID-19 Risk and Severity Is Increased in Vitamin D Deficiency. Front Public Health 8, 513.

Bleizgys, A. (2020). Vitamin D AND COVID-19: It is time to act. Int J Clin Pract, e13748.

Bleriot, C. and Ginhoux, F. (2019). Understanding the Heterogeneity of Resident Liver Macrophages. Front Immunol 10, 2694.

Cao, X., Lu, X. M., Tuo, X., Liu, J. Y., Zhang, Y. C., Song, L. N., Cheng, Z. Q., Yang, J. K. and Xin, Z. (2019). Angiotensin-converting enzyme 2 regulates endoplasmic reticulum stress and mitochondrial function to preserve skeletal muscle lipid metabolism. Lipids Health Dis 18, 207.

Cao, X., Yang, F., Shi, T., Yuan, M., Xin, Z., Xie, R., Li, S., Li, H. and Yang, J. K. (2016). Angiotensin-converting enzyme 2/angiotensin-(1-7)/Mas axis activates Akt signaling to ameliorate hepatic steatosis. Sci Rep 6, 21592.

Cao, X., Yang, F. Y., Xin, Z., Xie, R. R. and Yang, J. K. (2014). The ACE2/Ang-(1-7)/Mas axis can inhibit hepatic insulin resistance. Mol Cell Endocrinol 393, 30–8.

Chen, L., Marishta, A., Ellison, C. E. and Verzi, M. P. (2020a). Identification of Transcription Factors Regulating SARS-CoV-2 Entry Genes in the Intestine. Cell Mol Gastroenterol Hepatol.

Chen, L., Wang, L., Cheng, Q., Tu, Y. X., Yang, Z., Li, R. Z., Luo, Z. H. and Chen, Z. X. (2020b). Anti-masculinization induced by aromatase inhibitors in adult female zebrafish. BMC Genomics 21, 22.

Cheng, P. Y., Lin, C. C., Wu, C. S., Lu, Y. F., Lin, C. Y., Chung, C. C., Chu, C. Y., Huang, C. J., Tsai, C. Y., Korzh, S. et al. (2008). Zebrafish cdx1b regulates expression of downstream factors of Nodal signaling during early endoderm formation. Development 135, 941–52.

Cheng, Z. and Liu, Z. (2019). Renin-angiotensin system gene polymorphisms and colorectal cancer risk: a meta-analysis. J Renin Angiotensin Aldosterone Syst 20, 1470320319881932.

Chou, C. F., Loh, C. B., Foo, Y. K., Shen, S., Fielding, B. C., Tan, T. H., Khan, S., Wang, Y., Lim, S. G., Hong, W. et al. (2006). ACE2 orthologues in non-mammalian vertebrates (Danio, Gallus, Fugu, Tetraodon and Xenopus). Gene 377, 46–55.

Chua, R. L., Lukassen, S., Trump, S., Hennig, B. P., Wendisch, D., Pott, F., Debnath, O., Thurmann, L., Kurth, F., Volker, M. T. et al. (2020). COVID-19 severity correlates with airway epithelium-immune cell interactions identified by single-cell analysis. Nat Biotechnol 38, 970–979.

Coccheri, S. (2020). COVID-19: The crucial role of blood coagulation and fibrinolysis. Intern Emerg Med.

Dobin, A., Davis, C. A., Schlesinger, F., Drenkow, J., Zaleski, C., Jha, S., Batut, P., Chaisson, M. and Gingeras, T. R. (2013). STAR: ultrafast universal RNA-seq aligner. Bioinformatics 29, 15–21.

Domazet-Loso, T. and Tautz, D. (2010). A phylogenetically based transcriptome age index mirrors ontogenetic divergence patterns. Nature 468, 815–8.

Donmez, A., Rifaioglu, A. S., Acar, A., Dogan, T., Cetin-Atalay, R. and Atalay, V. (2020). iBioProVis: interactive visualization and analysis of compound bioactivity space. Bioinformatics 36, 4227–4230.

Duan, X., Han, Y., Yang, L., Nilsson-Payant, B. E., Wang, P., Zhang, T., Xiang, J., Xu, D., Wang, X., Uhl, S. et al. (2020). Identification of Drugs Blocking SARS-CoV-2 Infection using Human Pluripotent Stem Cell-derived Colonic Organoids. bioRxiv, 2020.05.02.073320.

Flores, M. V., Hall, C. J., Davidson, A. J., Singh, P. P., Mahagaonkar, A. A., Zon, L. I., Crosier, K. E. and Crosier, P. S. (2008). Intestinal differentiation in zebrafish requires Cdx1b, a functional equivalent of mammalian Cdx2. Gastroenterology 135, 1665–75.

Forlano, R., Mullish, B. H., Mukherjee, S. K., Nathwani, R., Harlow, C., Crook, P., Judge, R., Soubieres, A., Middleton, P., Daunt, A. et al. (2020). In-hospital mortality is associated with inflammatory response in NAFLD patients admitted for COVID-19. PLoS One 15, e0240400.

Forn-Cuni, G., Varela, M., Pereiro, P., Novoa, B. and Figueras, A. (2017). Conserved gene regulation during acute inflammation between zebrafish and mammals. Sci Rep 7, 41905.

Froehlicher, M., Liedtke, A., Groh, K., Lopez-Schier, H., Neuhauss, S. C., Segner, H. and Eggen, R. I. (2009). Estrogen receptor subtype beta2 is involved in neuromast development in zebrafish (Danio rerio) larvae. Dev Biol 330, 32–43.

Gallagher, P. E., Ferrario, C. M. and Tallant, E. A. (2008). MAP kinase/phosphatase pathway mediates the regulation of ACE2 by angiotensin peptides. Am J Physiol Cell Physiol 295, C1169–74.

Ganz, J., Melancon, E., Wilson, C., Amores, A., Batzel, P., Strader, M., Braasch, I., Diba, P., Kuhlman, J. A., Postlethwait, J. H. et al. (2019). Epigenetic factors Dnmt1 and Uhrf1 coordinate intestinal development. Dev Biol 455, 473–484.

Gautier, L., Cope, L., Bolstad, B. M. and Irizarry, R. A. (2004). affy--analysis of Affymetrix GeneChip data at the probe level. Bioinformatics 20, 307–15.

Gava, E., Samad-Zadeh, A., Zimpelmann, J., Bahramifarid, N., Kitten, G. T., Santos, R. A., Touyz, R. M. and Burns, K. D. (2009). Angiotensin-(1-7) activates a tyrosine phosphatase and inhibits glucose-induced signalling in proximal tubular cells. Nephrol Dial Transplant 24, 1766–73.

Gomes, M. C. and Mostowy, S. (2020). The Case for Modeling Human Infection in Zebrafish. Trends Microbiol 28, 10–18.

Gupte, M., Boustany-Kari, C. M., Bharadwaj, K., Police, S., Thatcher, S., Gong, M. C., English, V. L. and Cassis, L. A. (2008). ACE2 is expressed in mouse adipocytes and regulated by a high-fat diet. Am J Physiol Regul Integr Comp Physiol 295, R781–8.

Gwathmey, T. M., Pendergrass, K. D., Reid, S. D., Rose, J. C., Diz, D. I. and Chappell, M. C. (2010). Angiotensin-(1-7)-angiotensin-converting enzyme 2 attenuates reactive oxygen species formation to angiotensin II within the cell nucleus. Hypertension 55, 166–71.

Hamming, I., Cooper, M. E., Haagmans, B. L., Hooper, N. M., Korstanje, R., Osterhaus, A. D., Timens, W., Turner, A. J., Navis, G. and van Goor, H. (2007). The emerging role of ACE2 in physiology and disease. J Pathol 212, 1–11.

Heiden, T. C., Struble, C. A., Rise, M. L., Hessner, M. J., Hutz, R. J. and Carvan, M. J., 3rd. (2008). Molecular targets of 2,3,7,8-tetrachlorodibenzo-p-dioxin (TCDD) within the zebrafish ovary: insights into TCDD-induced endocrine disruption and reproductive toxicity. Reprod Toxicol 25, 47–57.

Hirano, T. and Murakami, M. (2020). COVID-19: A New Virus, but a Familiar Receptor and Cytokine Release Syndrome. Immunity 52, 731–733.

Holden, L. A. and Brown, K. H. (2018). Baseline mRNA expression differs widely between common laboratory strains of zebrafish. Sci Rep 8, 4780.

Hu, B., Chen, H., Liu, X., Zhang, C., Cole, G. J., Lee, J. A. and Chen, X. (2013). Transgenic overexpression of cdx1b induces metaplastic changes of gene expression in zebrafish esophageal squamous epithelium. Zebrafish 10, 218–27.

Huang, R., Zhu, L., Wang, J., Xue, L., Liu, L., Yan, X., Huang, S., Li, Y., Yan, X., Zhang, B. et al. (2020). Clinical features of COVID-19 patients with non-alcoholic fatty liver disease. Hepatol Commun.

Jacob, V., Chernyavskaya, Y., Chen, X., Tan, P. S., Kent, B., Hoshida, Y. and Sadler, K. C. (2015). DNA hypomethylation induces a DNA replication-associated cell cycle arrest to block hepatic outgrowth in uhrf1 mutant zebrafish embryos. Development 142, 510–21.

Jia, H. P., Look, D. C., Shi, L., Hickey, M., Pewe, L., Netland, J., Farzan, M., Wohlford-Lenane, C., Perlman, S. and McCray, P. B., Jr. (2005). ACE2 receptor expression and severe acute respiratory syndrome coronavirus infection depend on differentiation of human airway epithelia. J Virol 79, 14614–21.

Jia, J., Qin, J., Yuan, X., Liao, Z., Huang, J., Wang, B., Sun, C. and Li, W. (2019). Microarray and metabolome analysis of hepatic response to fasting and subsequent refeeding in zebrafish (Danio rerio). BMC Genomics 20, 919.

Jin, J. M., Bai, P., He, W., Wu, F., Liu, X. F., Han, D. M., Liu, S. and Yang, J. K. (2020). Gender Differences in Patients With COVID-19: Focus on Severity and Mortality. Front Public Health 8, 152.

Johnson, E., Vu, L. and Matarese, L. E. (2018). Bacteria, Bones, and Stones: Managing Complications of Short Bowel Syndrome. Nutr Clin Pract 33, 454–466.

Jothimani, D., Venugopal, R., Abedin, M. F., Kaliamoorthy, I. and Rela, M. (2020). COVID-19 and the liver. J Hepatol.

Kim, S. M., Kim, Y. G., Jeong, K. H., Lee, S. H., Lee, T. W., Ihm, C. G. and Moon, J. Y. (2012). Angiotensin II-induced mitochondrial Nox4 is a major endogenous source of oxidative stress in kidney tubular cells. PLoS One 7, e39739.

Kimmel, C. B., Ballard, W. W., Kimmel, S. R., Ullmann, B. and Schilling, T. F. (1995). Stages of embryonic development of the zebrafish. Dev Dyn 203, 253–310.

Koch, B. E. V., Yang, S., Lamers, G., Stougaard, J. and Spaink, H. P. (2018). Intestinal microbiome adjusts the innate immune setpoint during colonization through negative regulation of MyD88. Nat Commun 9, 4099.

Kruglikov, I. L., Shah, M. and Scherer, P. E. (2020). Obesity and diabetes as comorbidities for COVID-19: Underlying mechanisms and the role of viral-bacterial interactions. Elife 9.

Kryuchkova-Mostacci, N. and Robinson-Rechavi, M. (2017). A benchmark of gene expression tissue-specificity metrics. Brief Bioinform 18, 205–214.

Lee, I. T., Nakayama, T., Wu, C. T., Goltsev, Y., Jiang, S., Gall, P. A., Liao, C. K., Shih, L. C., Schurch, C. M., McIlwain, D. R. et al. (2020a). ACE2 localizes to the respiratory cilia and is not increased by ACE inhibitors or ARBs. Nat Commun 11, 5453.

Lee, I. T., Nakayama, T., Wu, C. T., Goltsev, Y., Jiang, S., Gall, P. A., Liao, C. K., Shih, L. C., Schurch, C. M., McIlwain, D. R. et al. (2020b). Robust ACE2 protein expression localizes to the motile cilia of the respiratory tract epithelia and is not increased by ACE inhibitors or angiotensin receptor blockers. medRxiv.

Lin, L., Jiang, X., Zhang, Z., Huang, S., Zhang, Z., Fang, Z., Gu, Z., Gao, L., Shi, H., Mai, L. et al. (2020). Gastrointestinal symptoms of 95 cases with SARS-CoV-2 infection. Gut 69, 997–1001.

Liu, W., Qiao, Q., Chen, Y., Wu, K. and Zhang, X. (2014). Microcystin-LR exposure to adult zebrafish (Danio rerio) leads to growth inhibition and immune dysfunction in F1 offspring, a parental transmission effect of toxicity. Aquat Toxicol 155, 360–7.

Logan, S. L., Thomas, J., Yan, J., Baker, R. P., Shields, D. S., Xavier, J. B., Hammer, B. K. and Parthasarathy, R. (2018). The Vibrio cholerae type VI secretion system can modulate host intestinal mechanics to displace gut bacterial symbionts. Proc Natl Acad Sci U S A 115, E3779–E3787.

Love, M. I., Huber, W. and Anders, S. (2014). Moderated estimation of fold change and dispersion for RNA-seq data with DESeq2. Genome Biol 15, 550.

Lu, H., Cui, Y., Jiang, L. and Ge, W. (2017). Functional Analysis of Nuclear Estrogen Receptors in Zebrafish Reproduction by Genome Editing Approach. Endocrinology 158, 2292–2308.

MacInnes, A. W., Amsterdam, A., Whittaker, C. A., Hopkins, N. and Lees, J. A. (2008). Loss of p53 synthesis in zebrafish tumors with ribosomal protein gene mutations. Proc Natl Acad Sci U S A 105, 10408–13.

Mahamid, M., Nseir, W., Khoury, T., Mahamid, B., Nubania, A., Sub-Laban, K., Schifter, J., Mari, A., Sbeit, W. and Goldin, E. (2020). Nonalcoholic fatty liver disease is associated with COVID-19 severity independently of metabolic syndrome: a retrospective case-control study. Eur J Gastroenterol Hepatol.

Marjoram, L., Alvers, A., Deerhake, M. E., Bagwell, J., Mankiewicz, J., Cocchiaro, J. L., Beerman, R. W., Willer, J., Sumigray, K. D., Katsanis, N. et al. (2015). Epigenetic control of intestinal barrier function and inflammation in zebrafish. Proc Natl Acad Sci U S A 112, 2770–5.

McCarthy, D. J., Chen, Y. and Smyth, G. K. (2012). Differential expression analysis of multifactor RNA-Seq experiments with respect to biological variation. Nucleic Acids Res 40, 4288–97.

Mickiewicz, K. M., Kawai, Y., Drage, L., Gomes, M. C., Davison, F., Pickard, R., Hall, J., Mostowy, S., Aldridge, P. D. and Errington, J. (2019). Possible role of L-form switching in recurrent urinary tract infection. Nat Commun 10, 4379.

Musavi, H., Abazari, O., Barartabar, Z., Kalaki-Jouybari, F., Hemmati-Dinarvand, M., Esmaeili, P. and Mahjoub, S. (2020). The benefits of Vitamin D in the COVID-19 pandemic: biochemical and immunological mechanisms. Arch Physiol Biochem, 1–9.

Mutanen, A., Barrett, M., Feng, Y., Lohi, J., Rabah, R., Teitelbaum, D. H. and Pakarinen, M. P. (2019). Short bowel mucosal morphology, proliferation and inflammation at first and repeat STEP procedures. J Pediatr Surg 54, 511–516.

Nehme, A., Zouein, F. A., Zayeri, Z. D. and Zibara, K. (2019). An Update on the Tissue Renin Angiotensin System and Its Role in Physiology and Pathology. J Cardiovasc Dev Dis 6.

Noris, M., Benigni, A. and Remuzzi, G. (2020). The case of complement activation in COVID-19 multiorgan impact. Kidney Int 98, 314–322.

Okuda, Y., Ogura, E., Kondoh, H. and Kamachi, Y. (2010). B1 SOX coordinate cell specification with patterning and morphogenesis in the early zebrafish embryo. PLoS Genet 6, e1000936.

Papatheodorou, I., Moreno, P., Manning, J., Fuentes, A. M., George, N., Fexova, S., Fonseca, N. A., Fullgrabe, A., Green, M., Huang, N. et al. (2020). Expression Atlas update: from tissues to single cells. Nucleic Acids Res 48, D77–D83.

Pawlotsky, J. M. (2020). COVID-19 and the liver-related deaths to come. Nat Rev Gastroenterol Hepatol 17, 523–525.

Pinter, M. and Jain, R. K. (2017). Targeting the renin-angiotensin system to improve cancer treatment: Implications for immunotherapy. Sci Transl Med 9.

Postlethwait, J. H., Farnsworth, D. R. and Miller, A. C. (2020). An intestinal cell type in zebrafish is the nexus for the SARS-CoV-2 receptor and the Renin-Angiotensin-Aldosterone System that contributes to COVID-19 comorbidities. bioRxiv, 2020.09.01.278366.

Rasha, F., Ramalingam, L., Gollahon, L., Rahman, R. L., Rahman, S. M., Menikdiwela, K. and Moustaid-Moussa, N. (2019). Mechanisms linking the renin-angiotensin system, obesity, and breast cancer. Endocr Relat Cancer 26, R653–R672.

Ritchie, M. E., Phipson, B., Wu, D., Hu, Y., Law, C. W., Shi, W. and Smyth, G. K. (2015). limma powers differential expression analyses for RNA-sequencing and microarray studies. Nucleic Acids Res 43, e47.

Robinson, M. W., Harmon, C. and O’Farrelly, C. (2016). Liver immunology and its role in inflammation and homeostasis. Cell Mol Immunol 13, 267–76.

Robison, B. D., Drew, R. E., Murdoch, G. K., Powell, M., Rodnick, K. J., Settles, M., Stone, D., Churchill, E., Hill, R. A., Papasani, M. R. et al. (2008). Sexual dimorphism in hepatic gene expression and the response to dietary carbohydrate manipulation in the zebrafish (Danio rerio). Comp Biochem Physiol Part D Genomics Proteomics 3, 141–54.

Rogers, E. D., Henry, T. B., Twiner, M. J., Gouffon, J. S., McPherson, J. T., Boyer, G. L., Sayler, G. S. and Wilhelm, S. W. (2011). Global gene expression profiling in larval zebrafish exposed to microcystin-LR and microcystis reveals endocrine disrupting effects of Cyanobacteria. Environ Sci Technol 45, 1962–9.

San, B., Aben, M., Elurbe, D. M., Voeltzke, K., Den Broeder, M. J., Rougeot, J., Legler, J. and Kamminga, L. M. (2018). Genetic and epigenetic regulation of zebrafish intestinal development. Epigenomes 2, 19.

Schall, K. A., Thornton, M. E., Isani, M., Holoyda, K. A., Hou, X., Lien, C. L., Grubbs, B. H. and Grikscheit, T. C. (2017). Short bowel syndrome results in increased gene expression associated with proliferation, inflammation, bile acid synthesis and immune system activation: RNA sequencing a zebrafish SBS model. BMC Genomics 18, 23.

Shannon, P., Markiel, A., Ozier, O., Baliga, N. S., Wang, J. T., Ramage, D., Amin, N., Schwikowski, B. and Ideker, T. (2003). Cytoscape: a software environment for integrated models of biomolecular interaction networks. Genome Res 13, 2498–504.

Sheahan, T., Morrison, T. E., Funkhouser, W., Uematsu, S., Akira, S., Baric, R. S. and Heise, M. T. (2008). MyD88 is required for protection from lethal infection with a mouse-adapted SARS-CoV. PLoS Pathog 4, e1000240.

Shehwana, H. and Konu, O. (2019). Comparative Transcriptomics Between Zebrafish and Mammals: A Roadmap for Discovery of Conserved and Unique Signaling Pathways in Physiology and Disease. Front Cell Dev Biol 7, 5.

Shi, T. T., Yang, F. Y., Liu, C., Cao, X., Lu, J., Zhang, X. L., Yuan, M. X., Chen, C. and Yang, J. K. (2018). Angiotensin-converting enzyme 2 regulates mitochondrial function in pancreatic beta-cells. Biochem Biophys Res Commun 495, 860–866.

Shoemaker, R., Tannock, L. R., Su, W., Gong, M., Gurley, S. B., Thatcher, S. E., Yiannikouris, F., Ensor, C. M. and Cassis, L. A. (2019). Adipocyte deficiency of ACE2 increases systolic blood pressures of obese female C57BL/6 mice. Biol Sex Differ 10, 45.

Silberg, D. G., Swain, G. P., Suh, E. R. and Traber, P. G. (2000). Cdx1 and cdx2 expression during intestinal development. Gastroenterology 119, 961–71.

Small, C. M., Carney, G. E., Mo, Q., Vannucci, M. and Jones, A. G. (2009). A microarray analysis of sex- and gonad-biased gene expression in the zebrafish: evidence for masculinization of the transcriptome. BMC Genomics 10, 579.

Song, L. N., Liu, J. Y., Shi, T. T., Zhang, Y. C., Xin, Z., Cao, X. and Yang, J. K. (2020). Angiotensin-(1-7), the product of ACE2 ameliorates NAFLD by acting through its receptor Mas to regulate hepatic mitochondrial function and glycolipid metabolism. FASEB J.

Song, R., Preston, G. and Yosypiv, I. V. (2012). Ontogeny of angiotensin-converting enzyme 2. Pediatr Res 71, 13–9.

Soni, K., Choudhary, A., Patowary, A., Singh, A. R., Bhatia, S., Sivasubbu, S., Chandrasekaran, S. and Pillai, B. (2013). miR-34 is maternally inherited in Drosophila melanogaster and Danio rerio. Nucleic Acids Res 41, 4470–80.

Stein, R. A. and Young, L. M. (2020). From ACE2 to COVID-19: A Multiorgan Endothelial Disease. Int J Infect Dis.

Stuckenholz, C., Lu, L., Thakur, P., Kaminski, N. and Bahary, N. (2009). FACS-assisted microarray profiling implicates novel genes and pathways in zebrafish gastrointestinal tract development. Gastroenterology 137, 1321–32.

Suarez-Farinas, M., Tokuyama, M., Wei, G., Huang, R., Livanos, A., Jha, D., Levescot, A., Irizar, H., Kosoy, R., Cording, S. et al. (2020). Intestinal inflammation modulates the expression of ACE2 and TMPRSS2 and potentially overlaps with the pathogenesis of SARS-CoV-2 related disease. Gastroenterology.

Szklarczyk, D., Franceschini, A., Wyder, S., Forslund, K., Heller, D., Huerta-Cepas, J., Simonovic, M., Roth, A., Santos, A., Tsafou, K. P. et al. (2015). STRING v10: protein-protein interaction networks, integrated over the tree of life. Nucleic Acids Res 43, D447–52.

Thakur, P. C., Davison, J. M., Stuckenholz, C., Lu, L. and Bahary, N. (2014). Dysregulated phosphatidylinositol signaling promotes endoplasmic-reticulum-stress-mediated intestinal mucosal injury and inflammation in zebrafish. Dis Model Mech 7, 93–106.

Vargas, M., Servillo, G. and Einav, S. (2020). Lopinavir/ritonavir for the treatment of SARS, MERS and COVID-19: a systematic review. Eur Rev Med Pharmacol Sci 24, 8592–8605.

Verstockt, B., Verstockt, S., Abdu Rahiman, S., Ke, B. J., Arnauts, K., Cleynen, I., Sabino, J., Ferrante, M., Matteoli, G. and Vermeire, S. (2020). Intestinal receptor of SARS-CoV-2 in inflamed IBD tissue seems downregulated by HNF4A in ileum and upregulated by interferon regulating factors in colon. J Crohns Colitis.

Wan, Y., Shang, J., Graham, R., Baric, R. S. and Li, F. (2020). Receptor Recognition by the Novel Coronavirus from Wuhan: an Analysis Based on Decade-Long Structural Studies of SARS Coronavirus. J Virol 94.

Wang, Y., Wang, Y., Luo, W., Huang, L., Xiao, J., Li, F., Qin, S., Song, X., Wu, Y., Zeng, Q. et al. (2020). A comprehensive investigation of the mRNA and protein level of ACE2, the putative receptor of SARS-CoV-2, in human tissues and blood cells. Int J Med Sci 17, 1522–1531.

White, R. J., Collins, J. E., Sealy, I. M., Wali, N., Dooley, C. M., Digby, Z., Stemple, D. L., Murphy, D. N., Billis, K., Hourlier, T. et al. (2017). A high-resolution mRNA expression time course of embryonic development in zebrafish. Elife 6.

Willis, A. R., Moore, C., Mazon-Moya, M., Krokowski, S., Lambert, C., Till, R., Mostowy, S. and Sockett, R. E. (2016). Injections of Predatory Bacteria Work Alongside Host Immune Cells to Treat Shigella Infection in Zebrafish Larvae. Curr Biol 26, 3343–3351.

Woolbright, B. L., Williams, C. D., Ni, H., Kumer, S. C., Schmitt, T., Kane, B. and Jaeschke, H. (2017). Microcystin-LR induced liver injury in mice and in primary human hepatocytes is caused by oncotic necrosis. Toxicon 125, 99–109.

Wu, K., Song, W., Zhang, Z. and Ge, W. (2020). Disruption of dmrt1 rescues the all-male phenotype of the cyp19a1a mutant in zebrafish - a novel insight into the roles of aromatase/estrogens in gonadal differentiation and early folliculogenesis. Development 147.

Yang, M., Ma, X., Xuan, X., Deng, H., Chen, Q. and Yuan, L. (2020). Liraglutide Attenuates Non-Alcoholic Fatty Liver Disease in Mice by Regulating the Local Renin-Angiotensin System. Front Pharmacol 11, 432.

Yao, T. T., Qian, J. D., Zhu, W. Y., Wang, Y. and Wang, G. Q. (2020). A systematic review of lopinavir therapy for SARS coronavirus and MERS coronavirus-A possible reference for coronavirus disease-19 treatment option. J Med Virol 92, 556–563.

Yaqoob, P. and Calder, P. C. (2007). Fatty acids and immune function: new insights into mechanisms. Br J Nutr 98 Suppl 1, S41–5.

Yildiz, G., Arslan-Ergul, A., Bagislar, S., Konu, O., Yuzugullu, H., Gursoy-Yuzugullu, O., Ozturk, N., Ozen, C., Ozdag, H., Erdal, E. et al. (2013). Genome-wide transcriptional reorganization associated with senescence-to-immortality switch during human hepatocellular carcinogenesis. PLoS One 8, e64016.

Yu, G., Wang, L. G., Han, Y. and He, Q. Y. (2012). clusterProfiler: an R package for comparing biological themes among gene clusters. OMICS 16, 284–7.

Yuan, J., Fan, D., Xue, Z., Qu, J. and Su, J. (2020). Co-Expression of Mitochondrial Genes and ACE2 in Cornea Involved in COVID-19. Invest Ophthalmol Vis Sci 61, 13.

Zaim, S., Chong, J. H., Sankaranarayanan, V. and Harky, A. (2020). COVID-19 and Multiorgan Response. Curr Probl Cardiol 45, 100618.

Zeituni, E. M., Wilson, M. H., Zheng, X., Iglesias, P. A., Sepanski, M. A., Siddiqi, M. A., Anderson, J. L., Zheng, Y. and Farber, S. A. (2016). Endoplasmic Reticulum Lipid Flux Influences Enterocyte Nuclear Morphology and Lipid-dependent Transcriptional Responses. J Biol Chem 291, 23804–23816.

Zhang, W., Xu, Y. Z., Liu, B., Wu, R., Yang, Y. Y., Xiao, X. Q. and Zhang, X. (2014). Pioglitazone upregulates angiotensin converting enzyme 2 expression in insulin-sensitive tissues in rats with high-fat diet-induced nonalcoholic steatohepatitis. ScientificWorldJournal 2014, 603409.

Ziegler, C. G. K., Allon, S. J., Nyquist, S. K., Mbano, I. M., Miao, V. N., Tzouanas, C. N., Cao, Y., Yousif, A. S., Bals, J., Hauser, B. M. et al. (2020). SARS-CoV-2 Receptor ACE2 Is an Interferon-Stimulated Gene in Human Airway Epithelial Cells and Is Detected in Specific Cell Subsets across Tissues. Cell 181, 1016–1035 e19.

