## Supplementary Information for "*ace2* expression is higher in intestines and liver while being tightly regulated in development and disease in zebrafish"

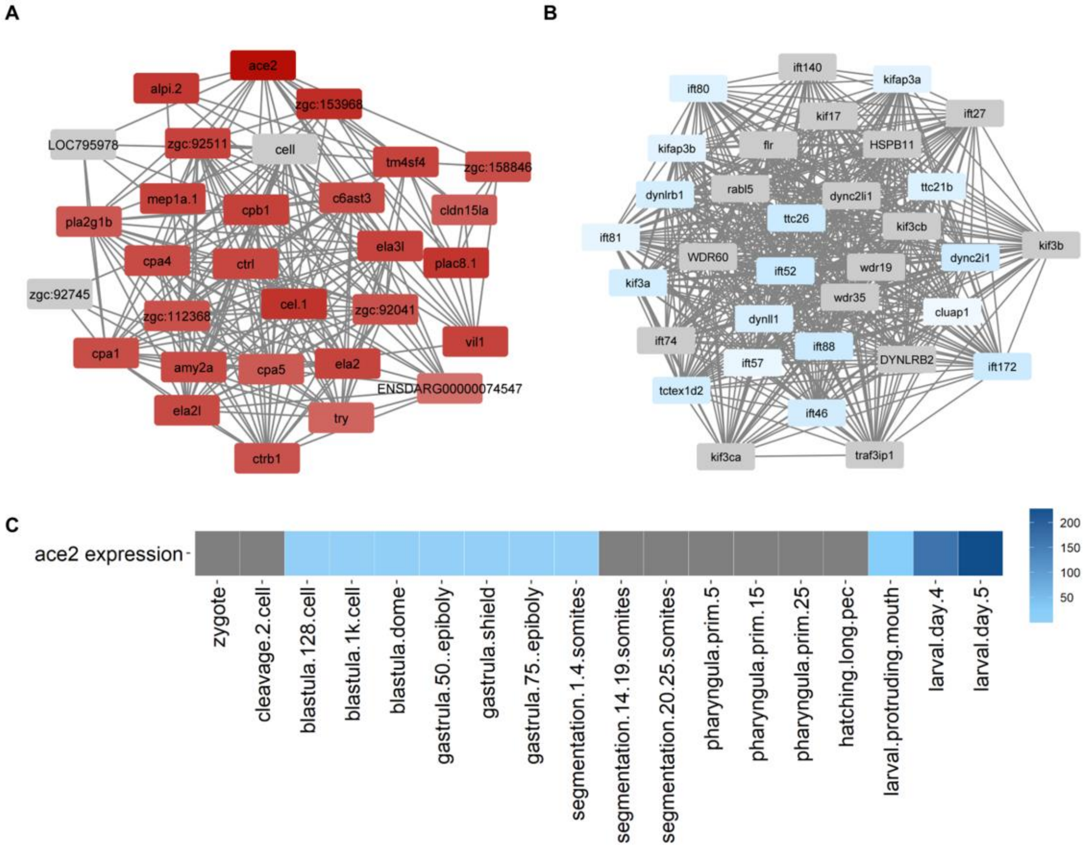


**Fig. S1.** Carboxypeptidase (A) and Intraflagellar (B) networks colored according to $r_{ace2}$ obtained from Affymetrix datasets. *ace2* expression levels during developmental stages from E-RAD475 dataset (C).

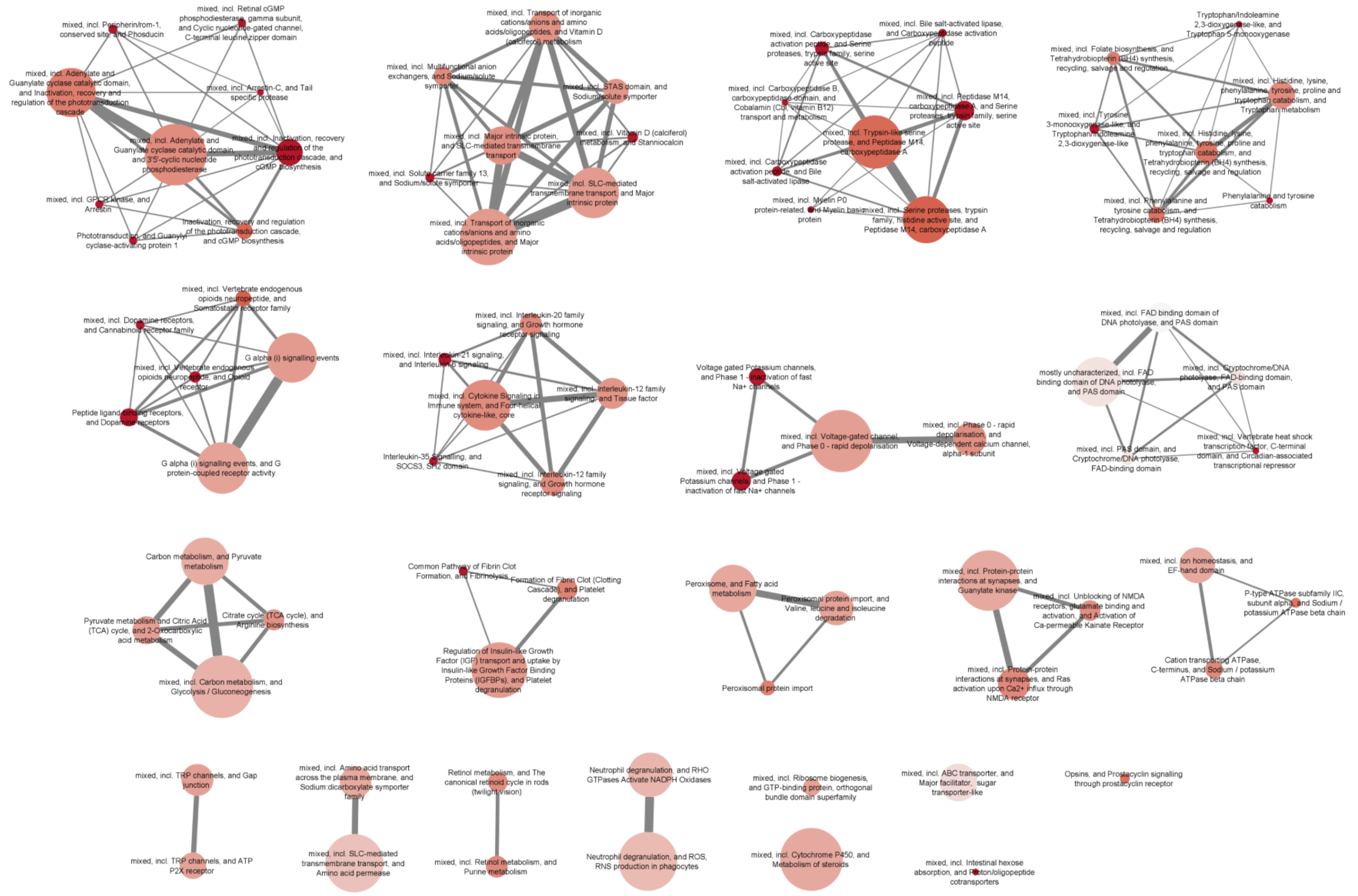


**Fig. S2**. Significant networks enriched in negative $r_{ace2}$ in development datasets (GSE24161 and GSE38575), where each node represented a local network, colored according to the mean STRING enrichment score and edge weights determined by the number of shared genes with other network it was connected to.

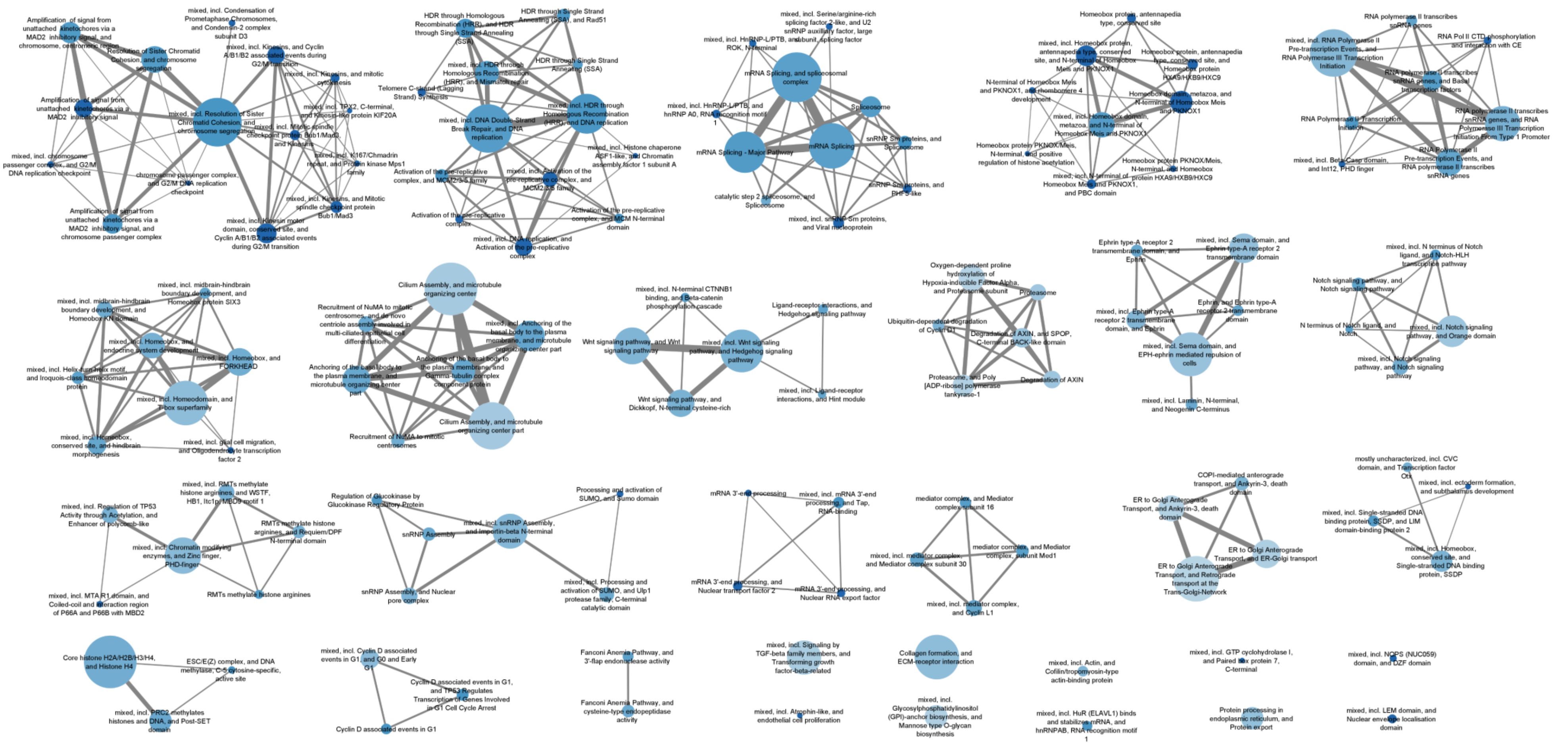


**Fig. S3.** Significant networks enriched in positive $r_{ace2}$ in development datasets (GSE24161 and GSE38575), where each node represented a local network, colored according to the mean STRING enrichment score and edge weights determined by the number of shared genes with other network it was connected to.

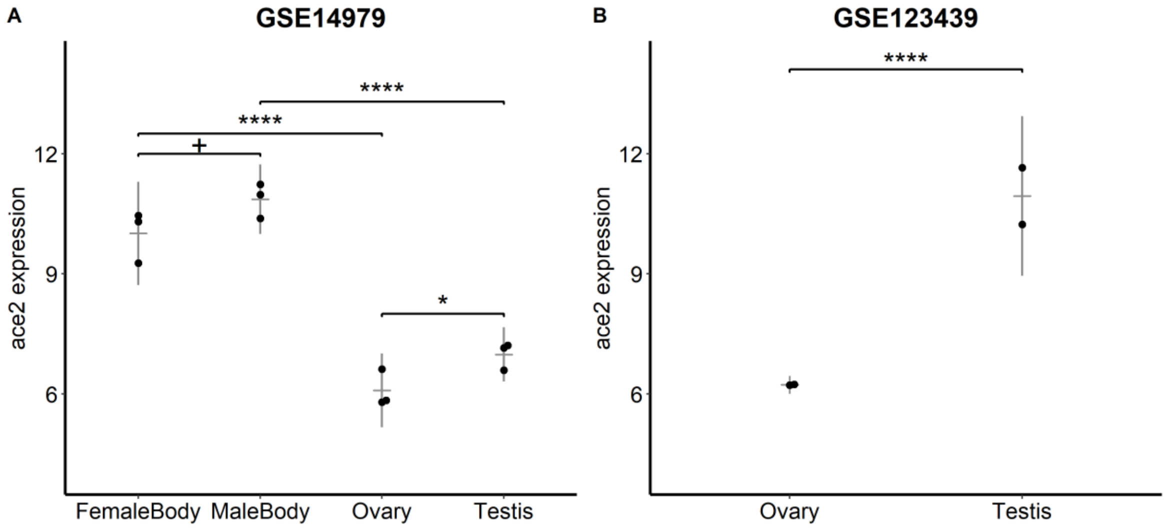


**Fig. S4.** ace2 expression from GSE14979 (A, Limma results were represented. +, adj. p value < 0.1; ****, adj. p-value < 0.0001) and GSE123439 (B, DESeq2 result was represented. ****, adj. p-value < 0.0001)

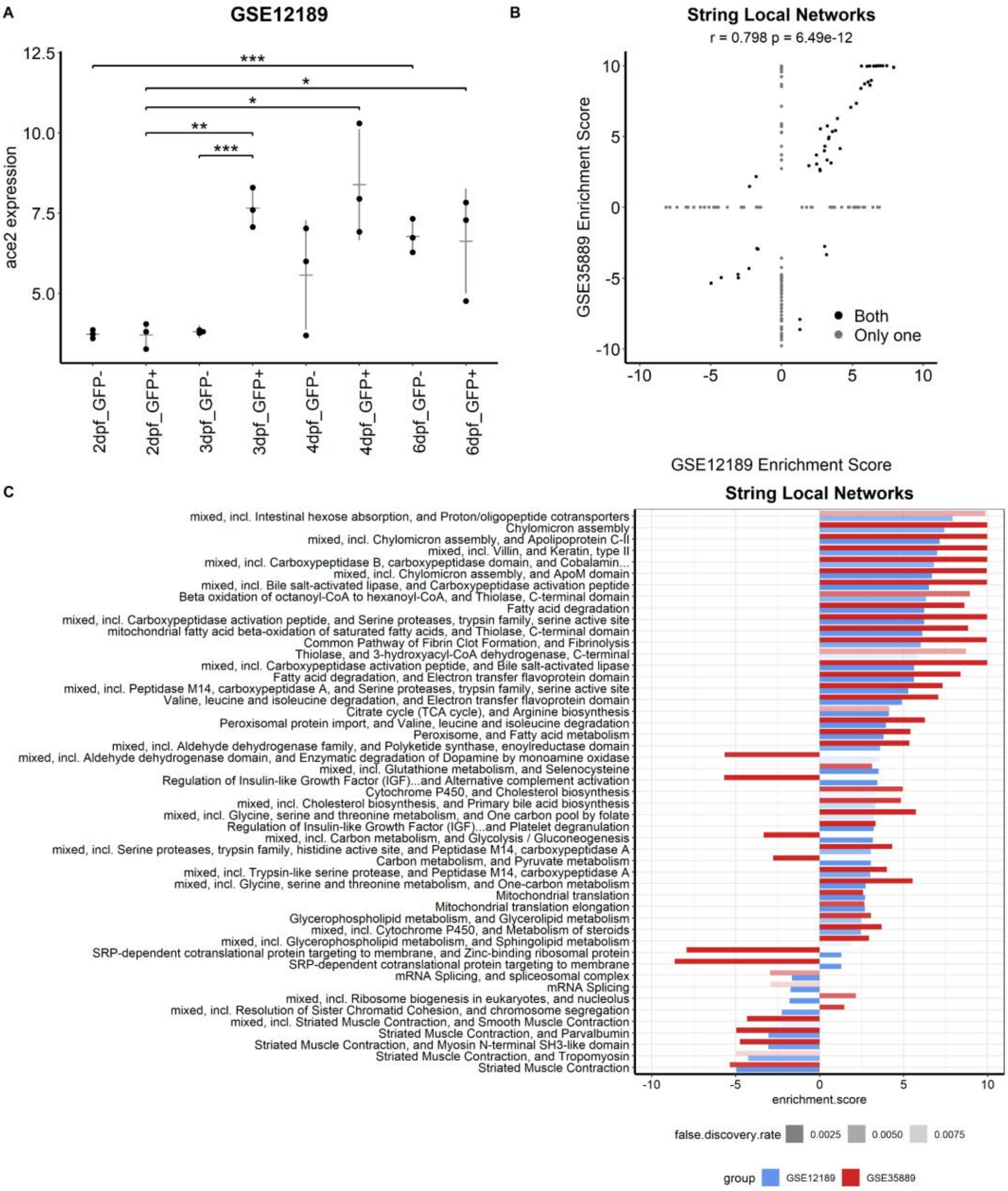


**Fig. S5**. ace2 expression from GSE12189 (A, Limma results were represented. *, adj. p-value < 0.05; **, adj. p-value < 0.01; ***, adj. p-value < 0.001; ****, adj. p-value < 0.0001). Correlation between significantly modulated STRING local networks with ace2 expression in GSE12189 and GSE35889 (B). Bar plots of significant STRING local network enrichment scores for GSE12189 and GSE35889 (C).

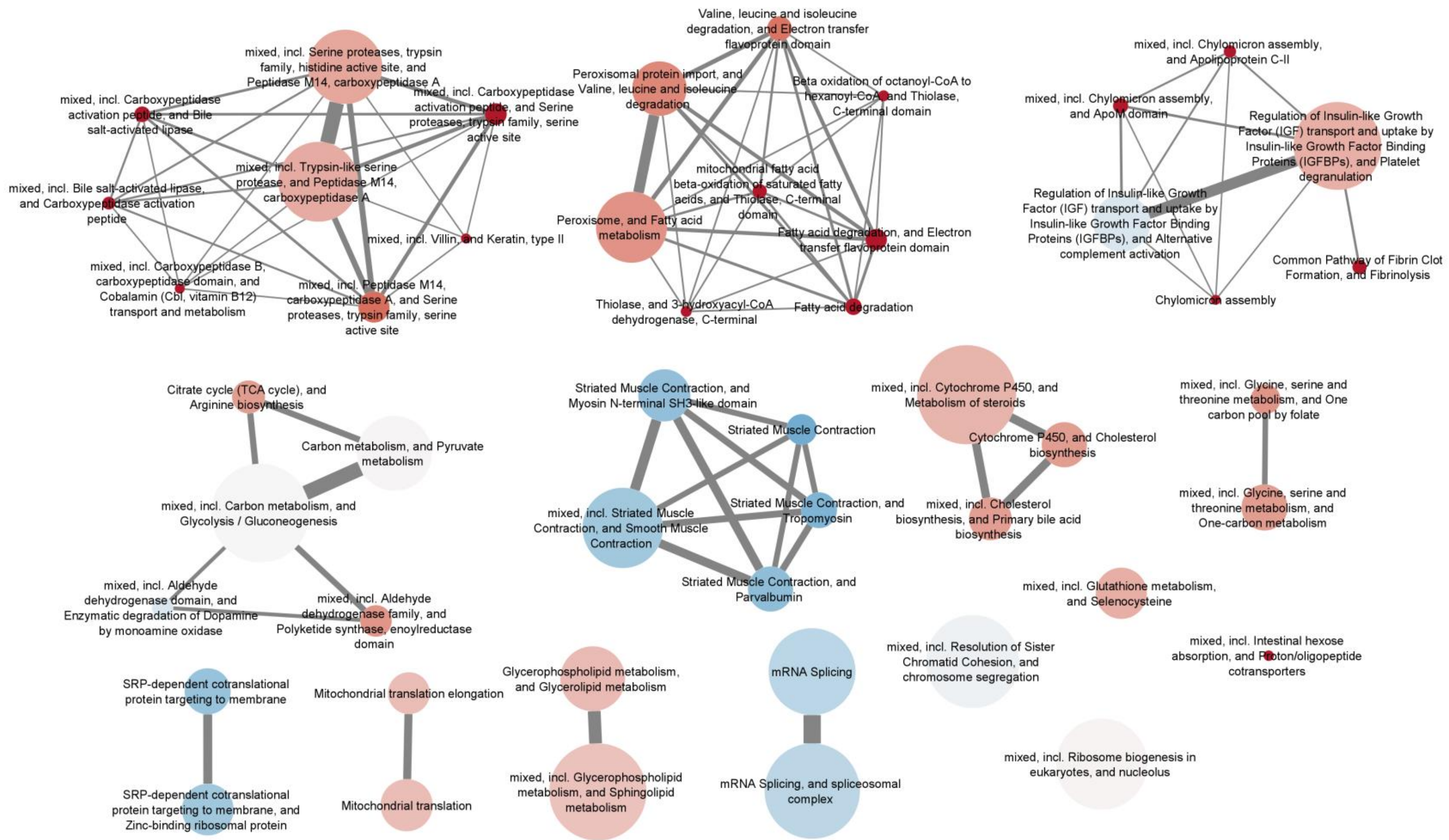


**Fig. S6.** Significant networks enriched in intestine related datasets (GSE35889 and GSE12189), where each node represented a local network, colored according to the mean STRING enrichment score and edge weights determined by the number of shared genes between networks.

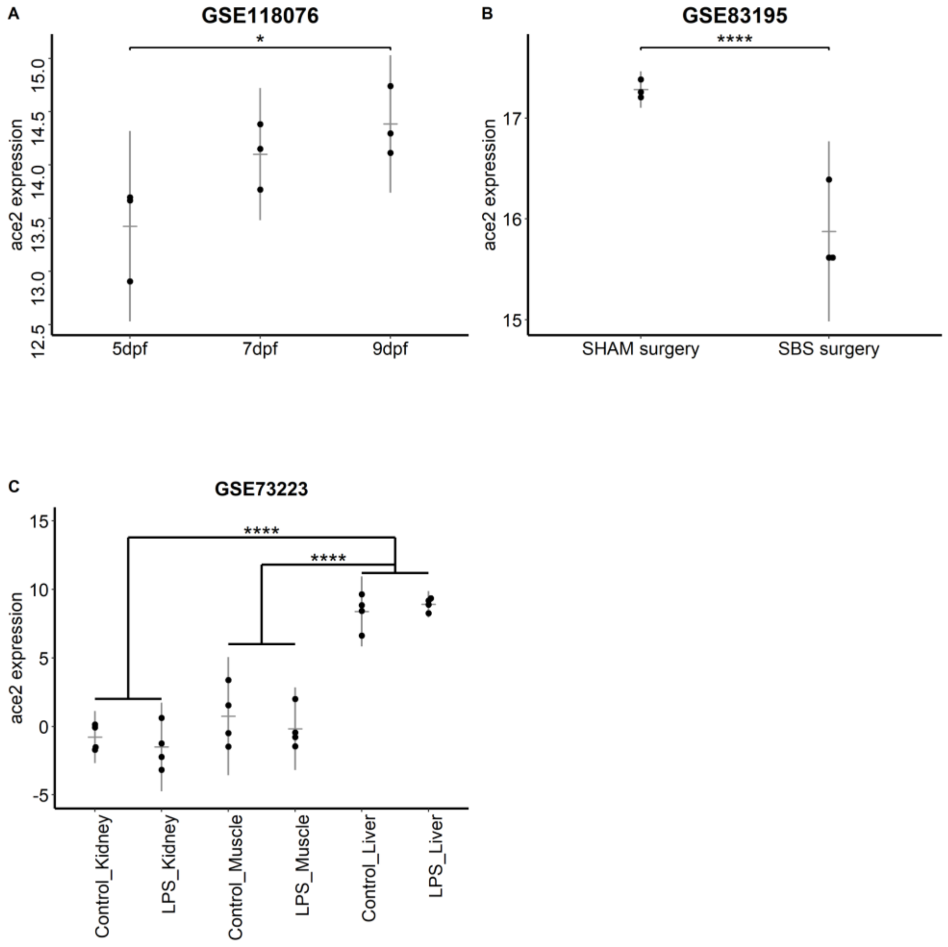


**Fig. S7.** *ace2* expression from GSE118076 (A, ANOVA results were represented. *, adj. p value < 0.05), GSE83195 (B, Student's t-test results were represented. ****, adj. p-value < 0.0001) and GSE73223 (C, Main factor analysis following two-way ANOVA results were represented. ****, adj. p-value < 0.0001).

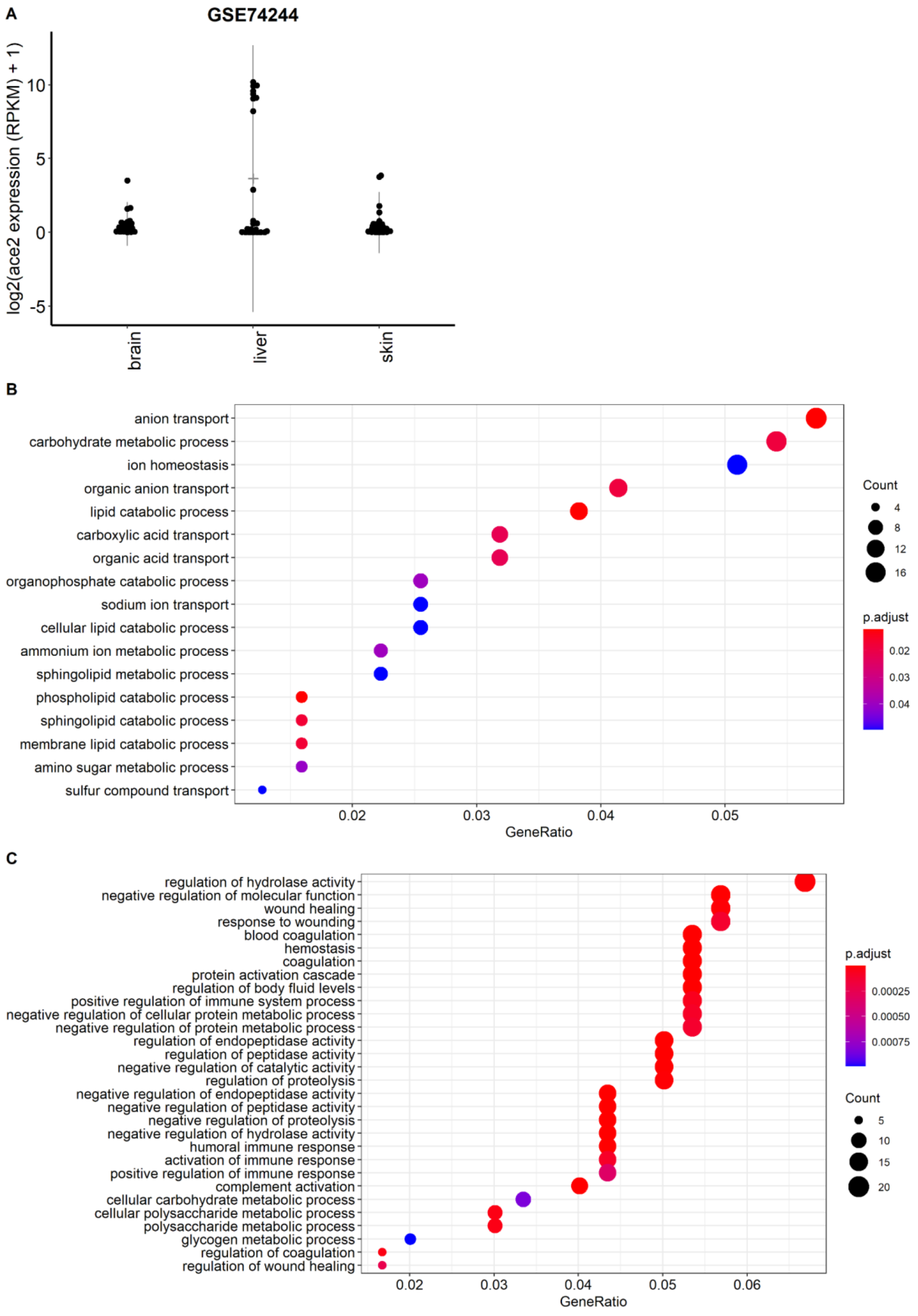


**Fig. S8.** *ace2* expression in liver. Tissue expression of *ace2* in brain, liver, and skin (A). GO term enrichment analysis results shown for positively correlated (B) and negatively correlated (C) genes.

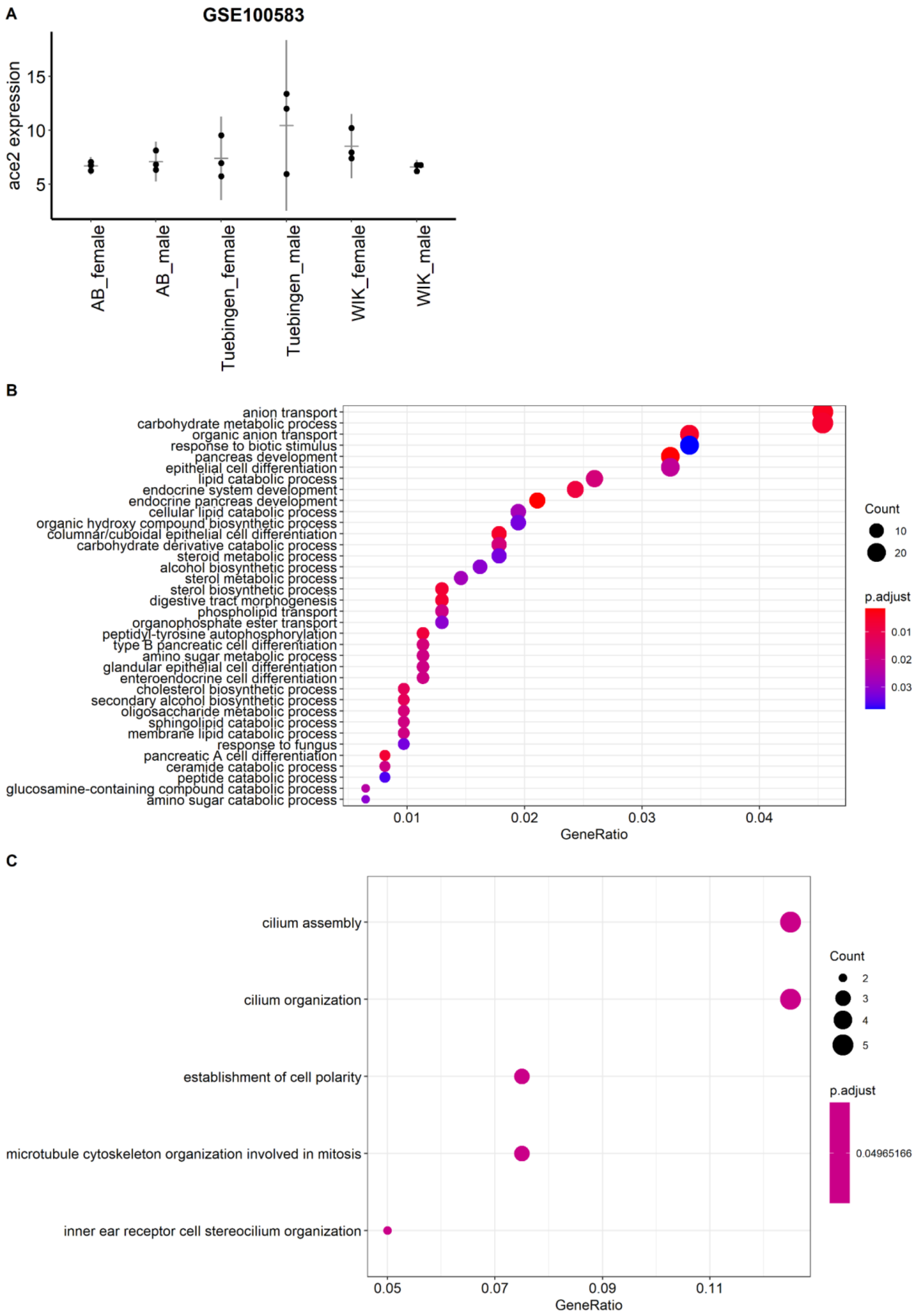


**Fig. S9.** *ace2* expression in GSE100583 (A). GO term enrichment analysis results shown for positively correlated (B) and negatively correlated (C) genes.

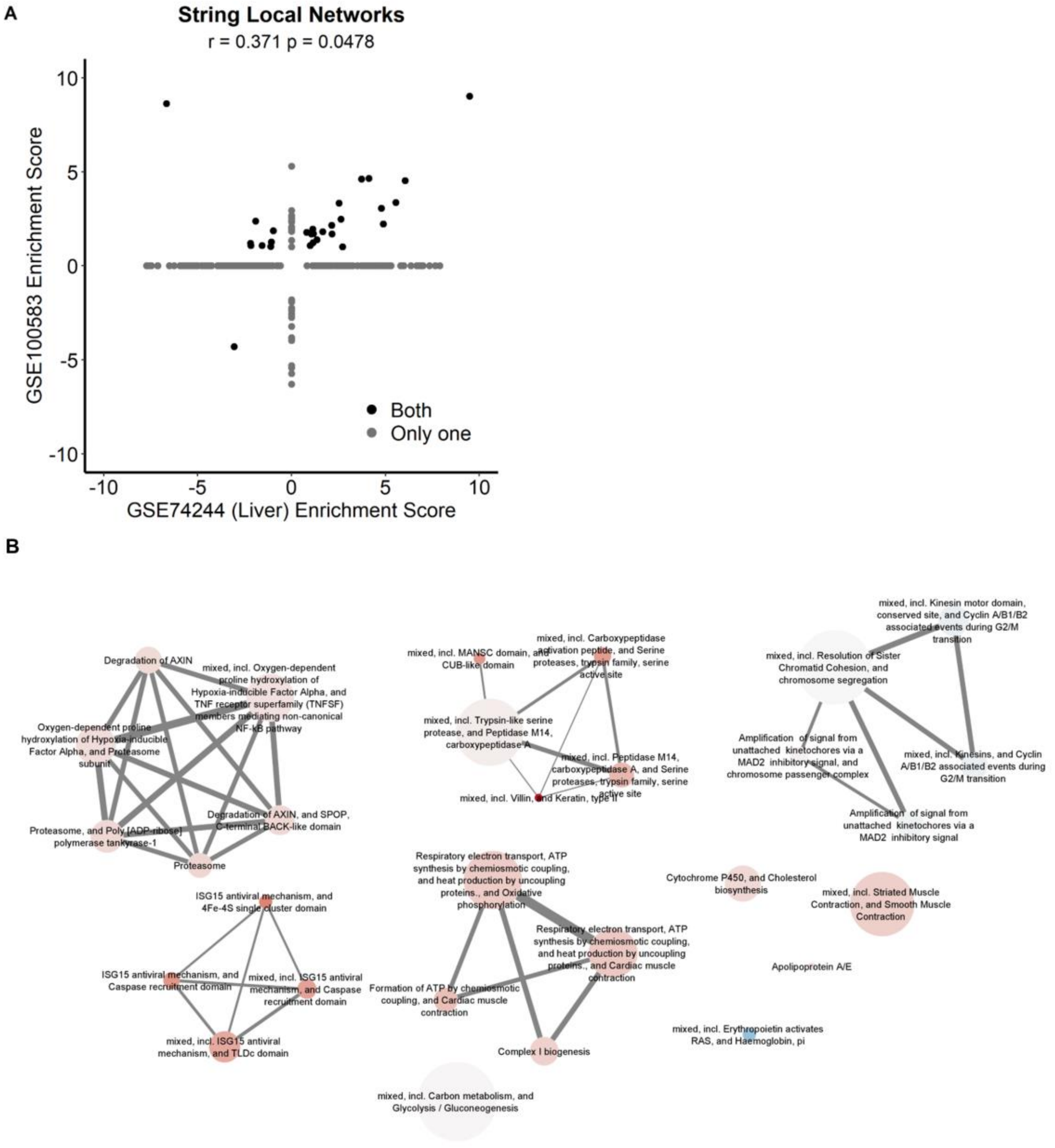


**Fig. S10.** Scatter plot of scores of local network enrichment analysis of GSE74244 and GSE100583 (A). Significant networks enriched in liver related datasets (GSE74244 and GSE100583), where each node represented a local network, colored according to the mean STRING enrichment score and edge weights indicate the number of shared genes between networks (B).

**Table S1.** Datasets used for tissue expression analysis

| GEO Accesion | Tissue Origin | Age/Time Point | Platform | PMID | Submission D | BioProject |
|---|---|---|---|---|---|---|
| **GSE74244** | Brain, liver, skin | 6 months | GPL14875 | 29382830 | Oct 21, 2015 | PRJNA299585 |
| **GSE46916** | Skin | 5 months | GPL15583 | 26620638 | May 14, 2013 | PRJNA203029 |
| **GSE73795** | Retina | 7-9 months | GPL14875 | NA | Oct 06, 2015 | PRJNA297904 |
| **GSE62221** | Brain, gill, heart, intestine, kidney, liver, muscle, spleen | 6 months | GPL14875 | 26227973 | Oct 09, 2014 | PRJNA263496 |
| **-** | Brain, gill, heart, intestine, kidney, muscle, spleen, bones, embryos, Unfertilized eggs, ovary, testis, liver | 5 months | GPL14875 | 27189481 | Jul 22,2014 | PRJNA255848 |

**Table S2.** Datasets used for analysis of *ace2* expression level.

| GEO Accesion | Tissue Origin | Age/Time Point | Number of groups | Total number of sample | Platform | PMID | Submission Date | BioProject |
|---|---|---|---|---|---|---|---|---|
| **GSE24616** | Whole organism | Egg - 1y9m | 72 | 147 | GPL6457 | 21150997 | Oct 11, 2010 | PRJNA132439 |
| **GSE82200** | Whole larvae | 5 dpf | 6 | 18 | GPL18413 | 30291253 | Jun 02, 2016 | PRJNA324288 |
| **GSE113241** | Kidney | Adults | 8 | 32 | GPL14664 | 30038625 | Apr 17, 2018 | PRJNA450536 |
| **GSE74244** | Brain, liver, skin | 6, 12, 24, 36, 42 m | 15 | 75 | GPL14875 | 29382830 | Oct 21, 2015 | PRJNA299585 |
| **GSE112272** | Liver | 3 months | 5 | 16 | GPL14664 | 31791229 | Mar 23, 2018 | PRJNA445444 |
| **GSE123439** | Gonad | 4 months | 8 | 16 | GPL23085 | 31910818 | Dec 06, 2018 | PRJNA508758 |
| **GSE118076** | Intestine | 5, 7, 9 dpf | 12 | 24 | GPL20828 | Doi: 10.3390/epigenomes2040019 | Aug 03, 2018 | PRJNA484346 |
| **GSE83195** | Intestine | Adult | 2 | 6 | GPL20828 | 28118819 | Jun 09, 2016 | PRJNA325275 |
| **GSE73223** | Liver, Kidney, Mus | Adult | 6 | 24 | GPL14664 | 28157230 | Sep 18, 2015 | PRJNA296395 |
| **GSE100583** | Liver | Adult | 6 | 18 | GPL14664 | 29555936 | Jun 27, 2017 | PRJNA392172 |

**Table S3.** Commonly Enriched Local Networks during larval development in GSE24616 (2 dpf – 8 dpf) and GSE38575 (2 dpf – 7 dpf). (ES; Enrichment Score, FDR; False Discovery Rate, ngenes; number of genes mapped)

| term.description | GSE24616 ngenes | GSE24616 ES | GSE24616 FDR | GSE38575 ngenes | GSE38575 ES | GSE38575 FDR |
|---|---|---|---|---|---|---|
| Activation of the pre-replicative complex | 11 | -6.88213 | 2.55E-06 | 12 | -8.73123 | 2.90E-05 |
| Activation of the pre-replicative complex, and MCM N-terminal domain | 15 | -6.76501 | 1.30E-06 | 17 | -7.18269 | 0.00038 |
| Activation of the pre-replicative complex, and MCM2/3/5 family | 25 | -6.28475 | 1.25E-09 | 24 | -7.52731 | 3.97E-06 |
| Amplification of signal from unattached kinetochores via a MAD2 inhibitory signal | 38 | -6.24383 | 1.23E-10 | 30 | -8.71292 | 8.36E-09 |
| Amplification of signal from unattached kinetochores via a MAD2 inhibitory signal, and chromosome passenger complex | 41 | -5.86703 | 5.28E-10 | 35 | -7.38952 | 6.10E-08 |
| Amplification of signal from unattached kinetochores via a MAD2 inhibitory signal, and chromosome, centromeric region | 43 | -5.78886 | 6.24E-10 | 36 | -7.11432 | 1.52E-07 |
| Anchoring of the basal body to the plasma membrane, and Gamma-tubulin complex component protein | 43 | -4.40511 | 5.77E-07 | 29 | -6.76796 | 0.00014 |
| Anchoring of the basal body to the plasma membrane, and microtubule organizing center part | 53 | -4.28205 | 2.86E-07 | 34 | -6.84965 | 1.23E-05 |
| Carbon metabolism, and Pyruvate metabolism | 110 | 2.18468 | 0.0055 | 104 | 4.72151 | 3.11E-08 |
| catalytic step 2 spliceosome, and Spliceosome | 21 | -4.10554 | 0.00012 | 19 | -6.53524 | 0.00062 |
| Cation transporting ATPase, C-terminus, and Sodium / potassium ATPase beta chain | 30 | 4.59523 | 0.0028 | 33 | 5.63456 | 0.00013 |
| chromosome passenger complex, and G2/M DNA replication checkpoint | 5 | -8.41381 | 1.74E-05 | 5 | -9.33092 | 0.0041 |
| Cilium Assembly, and microtubule organizing center | 107 | -2.62183 | 4.04E-05 | 81 | -3.77742 | 0.00042 |
| Cilium Assembly, and microtubule organizing center part | 102 | -2.70879 | 1.77E-05 | 77 | -3.84144 | 0.00029 |
| Citrate cycle (TCA cycle), and Arginine biosynthesis | 34 | 3.94408 | 0.0046 | 42 | 5.36883 | 0.00025 |
| Collagen formation, and ECM-receptor interaction | 67 | -2.42353 | 0.0017 | 69 | -5.72061 | 4.65E-08 |
| Common Pathway of Fibrin Clot Formation, and Fibrinolysis | 10 | 6.63416 | 0.0026 | 11 | 8.95068 | 1.12E-05 |
| COPI-mediated anterograde transport, and Ankyrin-3, death domain | 26 | -3.22132 | 0.00025 | 26 | -6.17509 | 0.0051 |

| | | | | | | |
|---|---|---|---|---|---|---|
| Core histone H2A/H2B/H3/H4, and Histone H4 | 58 | -3.36762 | 0.00011 | 49 | -6.99516 | 6.36E-09 |
| Cyclin D associated events in G1 | 15 | -6.24121 | 3.74E-06 | 15 | -7.43596 | 0.00051 |
| Cyclin D associated events in G1, and TP53 Regulates Transcription of Genes Involved in G1 Cell Cycle Arrest | 19 | -6.12116 | 1.30E-06 | 17 | -7.63969 | 8.40E-05 |
| Degradation of AXIN | 44 | -2.21628 | 0.0019 | 45 | -4.8216 | 3.08E-05 |
| Degradation of AXIN, and SPOP, C-terminal BACK-like domain | 49 | -2.28068 | 0.001 | 50 | -4.84977 | 1.67E-05 |
| Ephrin type-A receptor 2 transmembrane domain, and Ephrin | 18 | -4.1481 | 0.0045 | 16 | -7.07581 | 0.00085 |
| Ephrin, and Ephrin type-A receptor 2 transmembrane domain | 16 | -4.27619 | 0.0068 | 14 | -7.16737 | 0.0017 |
| ER to Golgi Anterograde Transport, and Ankyrin-3, death domain | 61 | -2.03008 | 0.0012 | 60 | -3.31241 | 0.005 |
| ER to Golgi Anterograde Transport, and ER-Golgi transport | 66 | -2.14039 | 0.00025 | 64 | -3.52457 | 0.0017 |
| ER to Golgi Anterograde Transport, and Retrograde transport at the Trans-Golgi-Network | 106 | -1.53168 | 4.69E-05 | 100 | -3.42509 | 2.46E-05 |
| ESC/E(Z) complex, and DNA methylase, C-5 cytosine-specific, active site | 10 | -5.67846 | 0.0019 | 11 | -8.16357 | 0.00082 |
| Fanconi Anemia Pathway, and 3'-flap endonuclease activity | 19 | -5.02364 | 9.74E-05 | 14 | -7.03722 | 0.0022 |
| Fanconi Anemia Pathway, and cysteine-type endopeptidase activity | 22 | -4.33325 | 0.00076 | 16 | -6.46031 | 0.0036 |
| Formation of Fibrin Clot (Clotting Cascade), and Platelet degranulation | 29 | 4.9926 | 0.00025 | 25 | 5.64429 | 0.0013 |
| G alpha (i) signalling events | 69 | 3.51638 | 4.25E-05 | 65 | 5.00267 | 2.22E-07 |
| G alpha (i) signalling events, and G protein-coupled receptor activity | 73 | 3.22618 | 7.10E-05 | 70 | 4.57902 | 7.25E-07 |
| HDR through Homologous Recombination (HRR), and HDR through Single Strand Annealing (SSA) | 40 | -4.94311 | 1.22E-09 | 30 | -8.62663 | 2.38E-10 |
| HDR through Single Strand Annealing (SSA) | 18 | -4.29609 | 0.0022 | 11 | -8.51588 | 0.00026 |
| HDR through Single Strand Annealing (SSA), and Rad51 | 27 | -4.78001 | 4.17E-06 | 17 | -8.53619 | 9.40E-07 |
| Homeobox domain, metazoa, and N-terminal of Homeobox Meis and PKNOX1 | 49 | -5.7919 | 6.91E-14 | 64 | -8.32432 | 2.53E-17 |

| | | | | | | |
|---|---|---|---|---|---|---|
| Homeobox protein PKNOX/Meis, N-terminal, and Homeobox protein HXA9/HXB9/HXC9 | 7 | -6.78813 | 0.0012 | 8 | -8.84018 | 0.0011 |
| Homeobox protein PKNOX/Meis, N-terminal, and positive regulation of histone acetylation | 5 | -6.88362 | 0.0065 | 4 | -9.57083 | 0.0016 |
| Homeobox protein, antennapedia type, conserved site | 14 | -6.76004 | 1.30E-06 | 20 | -9.13909 | 9.40E-07 |
| Homeobox protein, antennapedia type, conserved site, and Homeobox protein HXA9/HXB9/HXC9 | 18 | -6.6654 | 1.30E-06 | 26 | -9.06755 | 2.78E-10 |
| Inactivation, recovery and regulation of the phototransduction cascade, and cGMP biosynthesis | 25 | 5.80554 | 0.00027 | 17 | 7.47195 | 4.48E-05 |
| Interleukin-35 Signalling, and SOCS3, SH2 domain | 7 | 8.51266 | 0.00016 | 12 | 7.70124 | 0.00062 |
| Ligand-receptor interactions, and Hedgehog signaling pathway | 17 | -5.23738 | 8.64E-05 | 20 | -6.51218 | 0.00065 |
| mediator complex, and Mediator complex subunit 16 | 22 | -4.77245 | 0.00017 | 23 | -7.76856 | 6.13E-06 |
| mediator complex, and Mediator complex, subunit Med1 | 24 | -4.49499 | 0.00027 | 24 | -7.83687 | 2.64E-06 |
| mixed, incl. ABC transporter, and Major facilitator, sugar transporter-like | 46 | -1.93722 | 9.00E-04 | 42 | 4.58638 | 0.00017 |
| mixed, incl. Actin, and Cofilin/tropomyosin-type actin-binding protein | 21 | -3.3434 | 0.002 | 19 | -5.92536 | 0.0036 |
| mixed, incl. Activation of the pre-replicative complex, and MCM2/3/5 family | 32 | -6.54318 | 3.14E-13 | 29 | -7.74813 | 6.93E-08 |
| mixed, incl. Adenylate and Guanylate cyclase catalytic domain, and 3'5'-cyclic nucleotide phosphodiesterase | 86 | 4.50582 | 8.78E-11 | 68 | 5.40459 | 9.37E-10 |
| mixed, incl. Adenylate and Guanylate cyclase catalytic domain, and Inactivation, recovery and regulation of the phototransduction cascade | 73 | 5.08135 | 2.38E-11 | 57 | 6.09222 | 4.47E-10 |
| mixed, incl. Amino acid transport across the plasma membrane, and Sodium:dicarboxylate symporter family | 46 | 3.04425 | 0.0097 | 30 | 4.10088 | 0.0051 |
| mixed, incl. Anchoring of the basal body to the plasma membrane, and microtubule organizing center part | 57 | -4.44001 | 2.80E-08 | 36 | -7.00109 | 4.29E-06 |
| mixed, incl. Arrestin-C, and Tail specific protease | 5 | 8.8863 | 7.00E-04 | 4 | 9.47622 | 0.0036 |
| mixed, incl. Atrophin-like, and endothelial cell proliferation | 3 | -8.88839 | 0.00025 | 3 | -9.71355 | 0.0012 |
| mixed, incl. Beta-Casp domain, and Int12, PHD finger | 7 | -6.03745 | 0.009 | 7 | -9.12513 | 0.00065 |
| mixed, incl. Bile salt-activated lipase, and Carboxypeptidase activation peptide | 10 | 9.77226 | 1.30E-06 | 10 | 9.57946 | 9.40E-07 |

| | | | | | | |
|---|---|---|---|---|---|---|
| mixed, incl. Carbon metabolism, and Glycolysis / Gluconeogenesis | 145 | 2.26092 | 0.00091 | 140 | 4.19327 | 7.49E-09 |
| mixed, incl. Carboxypeptidase activation peptide, and Bile salt-activated lipase | 15 | 9.68624 | 2.26E-10 | 14 | 9.53099 | 9.40E-07 |
| mixed, incl. Carboxypeptidase activation peptide, and Serine proteases, trypsin family, serine active site | 26 | 9.66862 | 6.35E-19 | 18 | 8.88945 | 7.04E-09 |
| mixed, incl. Carboxypeptidase B, carboxypeptidase domain, and Cobalamin (Cbl, vitamin B12) transport and metabolism | 5 | 9.8143 | 1.30E-06 | 5 | 9.60833 | 0.00018 |
| mixed, incl. Chromatin modifying enzymes, and Zinc finger, PHD-finger | 77 | -3.54141 | 6.43E-07 | 62 | -5.0033 | 4.26E-08 |
| mixed, incl. chromosome passenger complex, and G2/M DNA replication checkpoint | 8 | -8.4004 | 1.30E-06 | 8 | -9.34114 | 3.46E-05 |
| mixed, incl. Condensation of Prometaphase Chromosomes, and Condensin-2 complex subunit D3 | 6 | -8.00576 | 2.49E-05 | 7 | -9.18162 | 5.00E-04 |
| mixed, incl. Cryptochrome/DNA photolyase, FAD-binding domain, and PAS domain | 22 | -4.23219 | 0.00021 | 26 | 6.24445 | 2.19E-05 |
| mixed, incl. Cyclin D associated events in G1, and G0 and Early G1 | 35 | -3.7307 | 0.00084 | 33 | -4.74518 | 0.0083 |
| mixed, incl. Cytochrome P450, and Metabolism of steroids | 138 | 3.48817 | 1.30E-06 | 136 | 3.75264 | 3.11E-06 |
| mixed, incl. Cytokine Signaling in Immune system, and Four-helical cytokine-like, core | 70 | 4.88252 | 2.65E-08 | 84 | 5.08088 | 2.36E-08 |
| mixed, incl. DNA Double-Strand Break Repair, and DNA replication | 114 | -5.11975 | 3.81E-24 | 95 | -7.45291 | 8.42E-23 |
| mixed, incl. DNA replication, and Activation of the pre-replicative complex | 38 | -6.64924 | 1.05E-16 | 36 | -8.03933 | 2.20E-10 |
| mixed, incl. Dopamine receptors, and Cannabinoid receptor family | 7 | 7.16009 | 0.0055 | 8 | 8.28466 | 0.0041 |
| mixed, incl. ectoderm formation, and subthalamus development | 6 | -6.3634 | 0.0093 | 5 | -9.45732 | 0.00091 |
| mixed, incl. Ephrin type-A receptor 2 transmembrane domain, and Ephrin | 28 | -3.5041 | 0.0016 | 29 | -5.54751 | 0.00094 |
| mixed, incl. FAD binding domain of DNA photolyase, and PAS domain | 25 | -3.78462 | 0.00014 | 47 | 3.67806 | 0.00025 |
| mixed, incl. Folate biosynthesis, and Tetrahydrobiopterin (BH4) synthesis, recycling, salvage and regulation | 19 | 5.68494 | 0.00031 | 23 | 5.59185 | 0.0055 |
| mixed, incl. glial cell migration, and Oligodendrocyte transcription factor 2 | 6 | -7.44243 | 0.00033 | 9 | -8.47615 | 0.0022 |
| mixed, incl. Glycosylphosphatidylinositol (GPI)-anchor biosynthesis, and Mannose type O-glycan biosynthesis | 48 | -1.6268 | 0.0021 | 45 | -3.91469 | 0.0065 |

| | | | | | | |
|---|---|---|---|---|---|---|
| mixed, incl. GPCR kinase, and Arrestin | 9 | 6.7026 | 0.0042 | 4 | 9.37727 | 0.0061 |
| mixed, incl. GTP cyclohydrolase I, and Paired box protein 7, C-terminal | 6 | -7.26634 | 7.00E-04 | 6 | -8.99865 | 0.0048 |
| mixed, incl. HDR through Homologous Recombination (HRR), and DNA replication | 94 | -5.40597 | 7.08E-24 | 76 | -8.03364 | 7.86E-24 |
| mixed, incl. HDR through Homologous Recombination (HRR), and Mismatch repair | 51 | -4.43159 | 5.20E-08 | 35 | -8.23408 | 2.74E-11 |
| mixed, incl. Helix-turn-helix motif, and Iroquois-class homeodomain protein | 28 | -4.43791 | 0.00033 | 32 | -7.50727 | 5.20E-07 |
| mixed, incl. Histidine, lysine, phenylalanine, tyrosine, proline and tryptophan catabolism, and Tetrahydrobiopterin (BH4) synthesis, recycling, salvage | 36 | 5.71653 | 8.17E-05 | 33 | 6.07592 | 5.66E-05 |
| mixed, incl. Histidine, lysine, phenylalanine, tyrosine, proline and tryptophan catabolism, and Tryptophan metabolism | 47 | 4.5081 | 0.00025 | 42 | 5.45507 | 0.00012 |
| mixed, incl. Histone chaperone ASF1-like, and Chromatin assembly factor 1 subunit A | 5 | -7.09218 | 0.0042 | 5 | -9.22471 | 0.0057 |
| mixed, incl. HnRNP-L/PTB, and hnRNP A0, RNA recognition motif 1 | 13 | -6.65859 | 1.30E-06 | 15 | -9.40732 | 9.40E-07 |
| mixed, incl. HnRNP-L/PTB, and ROK, N-terminal | 10 | -6.38036 | 0.00021 | 11 | -9.3858 | 9.40E-07 |
| mixed, incl. Homeobox domain, metazoa, and N-terminal of Homeobox Meis and PKNOX1 | 59 | -5.06992 | 2.73E-12 | 73 | -7.52643 | 2.59E-17 |
| mixed, incl. Homeobox protein, antennapedia type, conserved site, and N-terminal of Homeobox Meis and PKNOX1 | 46 | -5.95014 | 2.42E-14 | 58 | -8.5269 | 8.97E-17 |
| mixed, incl. Homeobox, and endocrine system development | 68 | -3.9819 | 2.86E-07 | 76 | -6.11561 | 6.17E-12 |
| mixed, incl. Homeobox, and FORKHEAD | 73 | -4.10076 | 1.47E-08 | 83 | -6.29737 | 8.63E-14 |
| mixed, incl. Homeobox, conserved site, and hindbrain morphogenesis | 45 | -4.44105 | 4.77E-06 | 53 | -6.8807 | 1.19E-09 |
| mixed, incl. Homeobox, conserved site, and Single-stranded DNA binding protein, SSDP | 45 | -3.7595 | 0.0031 | 45 | -5.81162 | 5.67E-05 |
| mixed, incl. Homeodomain, and T-box superfamily | 128 | -3.46647 | 1.24E-09 | 140 | -4.69124 | 2.12E-11 |
| mixed, incl. HuR (ELAVL1) binds and stabilizes mRNA, and hnRNPAB, RNA recognition motif 1 | 17 | -5.01586 | 0.00024 | 14 | -8.35534 | 4.01E-05 |
| mixed, incl. Inactivation, recovery and regulation of the phototransduction cascade, and cGMP biosynthesis | 47 | 6.98302 | 3.14E-13 | 36 | 8.31846 | 3.51E-13 |
| mixed, incl. Interleukin-12 family signaling, and Growth hormone receptor signaling | 29 | 4.35904 | 0.0016 | 52 | 5.38577 | 3.71E-05 |

| | | | | | | |
|---|---|---|---|---|---|---|
| mixed, incl. Interleukin-12 family signaling, and Tissue factor | 41 | 3.8957 | 0.00091 | 67 | 5.17513 | 7.10E-07 |
| mixed, incl. Interleukin-20 family signaling, and Growth hormone receptor signaling | 26 | 4.61184 | 8.00E-04 | 49 | 5.45919 | 4.65E-05 |
| mixed, incl. Interleukin-21 signaling, and Interleukin-6 signaling | 10 | 7.88752 | 5.53E-05 | 24 | 6.77289 | 0.00029 |
| mixed, incl. Intestinal hexose absorption, and Proton/oligopeptide cotransporters | 3 | 9.40227 | 0.0052 | 3 | 9.90973 | 3.14E-05 |
| mixed, incl. Ion homeostasis, and EF-hand domain | 58 | 3.36063 | 0.0039 | 49 | 3.86761 | 0.0053 |
| mixed, incl. K167/Chmadrin repeat, and Protein kinase Mps1 family | 4 | -8.64808 | 0.00011 | 4 | -9.47938 | 0.0036 |
| mixed, incl. Kinesin motor domain, conserved site, and Cyclin A/B1/B2 associated events during G2/M transition | 40 | -7.44137 | 2.27E-18 | 30 | -8.8752 | 1.28E-10 |
| mixed, incl. Kinesins, and Cyclin A/B1/B2 associated events during G2/M transition | 37 | -7.50244 | 1.25E-18 | 27 | -8.90547 | 1.33E-10 |
| mixed, incl. Kinesins, and mitotic cytokinesis | 12 | -8.50421 | 3.94E-08 | 12 | -9.42968 | 9.40E-07 |
| mixed, incl. Kinesins, and Mitotic spindle checkpoint protein Bub1/Mad3 | 22 | -8.25724 | 1.36E-12 | 19 | -9.35708 | 6.98E-09 |
| mixed, incl. Laminin, N-terminal, and Neogenin C-terminus | 14 | -4.74868 | 0.0028 | 15 | -6.30551 | 0.009 |
| mixed, incl. LEM domain, and Nuclear envelope localisation domain | 8 | -6.93241 | 0.00016 | 6 | -9.09236 | 0.0031 |
| mixed, incl. Ligand-receptor interactions, and Hint module | 14 | -5.2109 | 0.00072 | 15 | -6.31097 | 0.009 |
| mixed, incl. Major intrinsic protein, and SLC-mediated transmembrane transport | 67 | 3.8961 | 3.10E-05 | 69 | 5.59335 | 3.20E-08 |
| mixed, incl. mediator complex, and Cyclin L1 | 33 | -4.10104 | 0.00011 | 30 | -7.26078 | 1.20E-06 |
| mixed, incl. mediator complex, and Mediator complex subunit 30 | 28 | -4.26021 | 0.00034 | 27 | -7.29716 | 1.83E-06 |
| mixed, incl. midbrain-hindbrain boundary development, and Homeobox KN domain | 39 | -3.9793 | 0.00021 | 44 | -6.55436 | 4.08E-08 |
| mixed, incl. midbrain-hindbrain boundary development, and Homeobox protein SIX3 | 23 | -4.23512 | 0.0042 | 26 | -7.15052 | 3.08E-05 |
| mixed, incl. Mitotic spindle checkpoint protein Bub1/Mad3, and Kinesins | 15 | -8.27809 | 8.42E-09 | 14 | -9.46889 | 9.40E-07 |
| mixed, incl. mRNA 3'-end processing, and Tap, RNA-binding | 21 | -5.71667 | 1.27E-05 | 15 | -8.03332 | 8.40E-05 |

| | | | | | | |
|---|---|---|---|---|---|---|
| mixed, incl. MTA R1 domain, and Coiled-coil and interaction region of P66A and P66B with MBD2 | 8 | -5.89801 | 0.0042 | 6 | -9.48643 | 7.54E-05 |
| mixed, incl. Multifunctional anion exchangers, and Sodium/solute symporter | 27 | 4.10756 | 0.008 | 25 | 5.80871 | 0.00085 |
| mixed, incl. Myelin P0 protein-related, and Myelin basic protein | 3 | 9.28383 | 0.0062 | 4 | 9.7763 | 1.00E-04 |
| mixed, incl. N-terminal CTNNB1 binding, and Beta-catenin phosphorylation cascade | 15 | -4.56302 | 0.0035 | 16 | -7.77387 | 0.00012 |
| mixed, incl. N-terminal of Homeobox Meis and PKNOX1, and PBC domain | 16 | -6.49747 | 1.30E-06 | 17 | -8.9496 | 9.40E-07 |
| mixed, incl. N terminus of Notch ligand, and Notch-HLH transcription pathway | 15 | -5.29744 | 0.00031 | 17 | -7.39135 | 2.00E-04 |
| mixed, incl. NOPS (NUC059) domain, and DZF domain | 9 | -6.06516 | 0.0011 | 9 | -9.3745 | 4.82E-06 |
| mixed, incl. Notch signaling pathway, and Notch signaling pathway | 24 | -4.1193 | 0.0013 | 26 | -6.11748 | 4.06E-05 |
| mixed, incl. Notch signaling pathway, and Orange domain | 55 | -2.76633 | 0.0065 | 58 | -4.21576 | 2.00E-04 |
| mixed, incl. PAS domain, and Cryptochrome/DNA photolyase, FAD-binding domain | 19 | -4.15707 | 0.00018 | 23 | 5.86576 | 0.00039 |
| mixed, incl. Peptidase M14, carboxypeptidase A, and Serine proteases, trypsin family, serine active site | 36 | 8.5372 | 2.32E-17 | 26 | 8.53942 | 8.98E-11 |
| mixed, incl. Peripherin/rom-1, conserved site, and Phosducin | 12 | 8.84654 | 1.30E-06 | 9 | 9.41772 | 9.40E-07 |
| mixed, incl. Phase 0 - rapid depolarisation, and Voltage-dependent calcium channel, alpha-1 subunit | 44 | 3.97015 | 5.80E-05 | 34 | 5.5939 | 7.24E-06 |
| mixed, incl. Phenylalanine and tyrosine catabolism, and Tetrahydrobiopterin (BH4) synthesis, recycling, salvage and regulation | 28 | 6.09637 | 0.00038 | 28 | 5.96398 | 0.00025 |
| mixed, incl. PRC2 methylates histones and DNA, and Post-SET domain | 39 | -4.0143 | 0.00014 | 37 | -7.13763 | 1.13E-06 |
| mixed, incl. Processing and activation of SUMO, and Ulp1 protease family, C-terminal catalytic domain | 10 | -5.20874 | 0.0049 | 20 | -5.71087 | 0.0066 |
| mixed, incl. Protein-protein interactions at synapses, and Guanylate kinase | 75 | 2.81477 | 0.0066 | 74 | 4.45955 | 8.19E-05 |
| mixed, incl. Protein-protein interactions at synapses, and Ras activation upon Ca2+ influx through NMDA receptor | 37 | 3.80636 | 0.0039 | 40 | 6.44438 | 9.40E-07 |
| mixed, incl. Regulation of TP53 Activity through Acetylation, and Enhancer of polycomb-like | 35 | -4.20636 | 8.64E-05 | 27 | -5.51796 | 0.00016 |
| mixed, incl. Resolution of Sister Chromatid Cohesion, and chromosome segregation | 109 | -6.09788 | 5.12E-29 | 87 | -7.50703 | 4.82E-23 |

| | | | | | | |
|---|---|---|---|---|---|---|
| mixed, incl. Retinal cGMP phosphodiesterase, gamma subunit, and Cyclic nucleotide-gated channel, C-terminal leucine zipper domain | 5 | 8.32083 | 0.0044 | 6 | 8.6341 | 0.0082 |
| mixed, incl. Retinol metabolism, and Purine metabolism | 39 | 4.64503 | 2.72E-05 | 39 | 5.43102 | 1.72E-05 |
| mixed, incl. Ribosome biogenesis, and GTP-binding protein, orthogonal bundle domain superfamily | 34 | 3.34344 | 0.0025 | 27 | 4.64481 | 0.00075 |
| mixed, incl. RMTs methylate histone arginines, and WSTF, HB1, Itc1p, MBD9 motif 1 | 26 | -4.18695 | 0.00058 | 22 | -5.79712 | 1.63E-05 |
| mixed, incl. RNA Polymerase II Pre-transcription Events, and RNA Polymerase III Transcription Initiation | 118 | -3.22461 | 1.09E-09 | 107 | -5.5623 | 1.57E-09 |
| mixed, incl. Sema domain, and EPH-ephrin mediated repulsion of cells | 72 | -2.9083 | 0.00011 | 74 | -4.45528 | 1.67E-05 |
| mixed, incl. Sema domain, and Ephrin type-A receptor 2 transmembrane domain | 53 | -2.91507 | 0.00015 | 53 | -4.86087 | 2.72E-05 |
| mixed, incl. Serine proteases, trypsin family, histidine active site, and Peptidase M14, carboxypeptidase A | 71 | 6.09553 | 2.46E-12 | 40 | 7.79793 | 7.96E-10 |
| mixed, incl. Serine/arginine-rich splicing factor 2-like, and U2 snRNP auxilliary factor, large subunit, splicing factor | 10 | -5.71864 | 0.0017 | 8 | -9.02501 | 0.00044 |
| mixed, incl. Signaling by TGF-beta family members, and Transforming growth factor-beta-related | 71 | -2.37062 | 0.0029 | 78 | -3.71193 | 0.00091 |
| mixed, incl. Single-stranded DNA binding protein, SSDP, and LIM domain-binding protein 2 | 18 | -4.35944 | 0.0018 | 12 | -8.40138 | 2.00E-04 |
| mixed, incl. SLC-mediated transmembrane transport, and Amino acid permease | 92 | 2.0981 | 0.0087 | 80 | 2.84991 | 0.0041 |
| mixed, incl. SLC-mediated transmembrane transport, and Major intrinsic protein | 80 | 3.44819 | 8.27E-05 | 74 | 5.08001 | 7.11E-07 |
| mixed, incl. snRNP Assembly, and Importin-beta N-terminal domain | 76 | -3.43292 | 6.52E-07 | 68 | -5.43855 | 1.99E-06 |
| mixed, incl. snRNP Sm proteins, and Viral nucleoprotein | 23 | -5.72689 | 1.19E-06 | 22 | -8.62962 | 7.45E-08 |
| mixed, incl. Solute carrier family 13, and Sodium/solute symporter | 12 | 6.60029 | 0.00074 | 10 | 8.6695 | 0.00019 |
| mixed, incl. STAS domain, and Sodium/solute symporter | 37 | 3.6497 | 0.0085 | 35 | 5.47145 | 0.00013 |
| mixed, incl. TPX2, C-terminal, and Kinesin-like protein KIF20A | 3 | -8.54451 | 0.0017 | 3 | -9.57411 | 0.0088 |
| mixed, incl. Transport of inorganic cations/anions and amino acids/oligopeptides, and Major intrinsic protein | 89 | 3.36932 | 1.00E-04 | 81 | 5.21864 | 2.86E-08 |
| mixed, incl. Transport of inorganic cations/anions and amino acids/oligopeptides, and Vitamin D (calciferol) metabolism | 52 | 3.68239 | 0.0012 | 53 | 5.62868 | 1.00E-06 |

| | | | | | | |
|---|---|---|---|---|---|---|
| mixed, incl. TRP channels, and ATP P2X receptor | 32 | 3.58095 | 0.0096 | 32 | 4.21096 | 0.0046 |
| mixed, incl. TRP channels, and Gap junction | 39 | 3.66231 | 0.0034 | 36 | 4.53131 | 0.0018 |
| mixed, incl. Trypsin-like serine protease, and Peptidase M14, carboxypeptidase A | 81 | 5.49537 | 5.58E-11 | 46 | 7.61526 | 2.64E-10 |
| mixed, incl. Tyrosine 3-monooxygenase-like, and Tryptophan/Indoleamine 2,3-dioxygenase-like | 11 | 6.96434 | 6.00E-04 | 13 | 8.44235 | 2.12E-05 |
| mixed, incl. Unblocking of NMDA receptors, glutamate binding and activation, and Activation of Ca-permeable Kainate Receptor | 26 | 4.08015 | 0.0094 | 23 | 5.62052 | 2.00E-04 |
| mixed, incl. Vertebrate endogenous opioids neuropeptide, and Opioid receptor | 14 | 6.34148 | 5.00E-04 | 15 | 8.45598 | 2.58E-06 |
| mixed, incl. Vertebrate endogenous opioids neuropeptide, and Somatostatin receptor family | 23 | 6.09611 | 0.00011 | 20 | 7.89518 | 9.40E-07 |
| mixed, incl. Vertebrate heat shock transcription factor, C-terminal domain, and Circadian-associated transcriptional repressor | 4 | 9.54614 | 0.00012 | 5 | 9.80242 | 4.11E-06 |
| mixed, incl. Vitamin D (calciferol) metabolism, and Stanniocalcin | 12 | 6.33843 | 0.0015 | 15 | 8.05943 | 2.45E-05 |
| mixed, incl. Voltage-gated channel, and Phase 0 - rapid depolarisation | 76 | 3.2906 | 3.96E-06 | 56 | 4.87421 | 3.68E-07 |
| mixed, incl. Voltage gated Potassium channels, and Phase 1 - inactivation of fast Na+ channels | 14 | 6.30193 | 0.00057 | 10 | 8.30811 | 0.00051 |
| mixed, incl. Wnt signaling pathway, and Hedgehog signaling pathway | 89 | -3.40867 | 4.09E-07 | 96 | -6.28422 | 2.40E-13 |
| mostly uncharacterized, incl. CVC domain, and Transcription factor Otx | 20 | -4.03349 | 0.0049 | 22 | -6.11967 | 0.0031 |
| mostly uncharacterized, incl. FAD binding domain of DNA photolyase, and PAS domain | 58 | -2.13165 | 0.0054 | 68 | 4.00232 | 9.83E-05 |
| mRNA 3'-end processing | 10 | -7.01103 | 7.96E-06 | 8 | -8.85744 | 0.00097 |
| mRNA 3'-end processing, and Nuclear RNA export factor | 13 | -6.25976 | 1.64E-05 | 9 | -8.39699 | 0.0024 |
| mRNA 3'-end processing, and Nuclear transport factor 2 | 17 | -6.33823 | 1.30E-06 | 12 | -8.58527 | 6.67E-05 |
| mRNA Splicing | 132 | -4.92702 | 2.31E-28 | 125 | -7.39675 | 2.98E-25 |
| mRNA Splicing - Major Pathway | 120 | -5.08491 | 1.49E-27 | 113 | -7.73157 | 6.55E-26 |
| mRNA Splicing, and spliceosomal complex | 140 | -4.83832 | 5.44E-29 | 131 | -7.36849 | 5.39E-25 |

| | | | | | | |
|---|---|---|---|---|---|---|
| N-terminal of Homeobox Meis and PKNOX1, and rhombomere 4 development | 8 | -6.40463 | 0.0012 | 7 | -9.36982 | 7.54E-05 |
| N terminus of Notch ligand, and Notch | 12 | -5.54278 | 0.00072 | 13 | -6.92282 | 0.0038 |
| Neutrophil degranulation, and RHO GTPases Activate NADPH Oxidases | 72 | 2.47174 | 0.00056 | 64 | 3.59907 | 0.0027 |
| Neutrophil degranulation, and ROS, RNS production in phagocytes | 101 | 2.22428 | 0.00019 | 87 | 3.37072 | 0.00037 |
| Notch signaling pathway, and Notch signaling pathway | 21 | -4.93405 | 0.00091 | 25 | -6.73492 | 1.70E-05 |
| Opsins, and Prostacyclin signalling through prostacyclin receptor | 11 | 6.32463 | 0.003 | 15 | 6.03128 | 0.0068 |
| Oxygen-dependent proline hydroxylation of Hypoxia-inducible Factor Alpha, and Proteasome subunit | 65 | -1.64278 | 0.0065 | 69 | -3.56205 | 0.00016 |
| P-type ATPase subfamily IIC, subunit alpha, and Sodium / potassium ATPase beta chain | 16 | 5.14032 | 0.0071 | 16 | 6.89128 | 0.00041 |
| Peptide ligand-binding receptors, and Dopamine receptors | 26 | 6.15556 | 4.73E-05 | 24 | 8.06436 | 1.57E-06 |
| Peroxisomal protein import | 23 | 5.00713 | 0.0084 | 16 | 5.95494 | 0.0043 |
| Peroxisomal protein import, and Valine, leucine and isoleucine degradation | 76 | 3.39497 | 2.12E-05 | 61 | 4.39526 | 1.56E-05 |
| Peroxisome, and Fatty acid metabolism | 100 | 3.53175 | 7.17E-07 | 77 | 3.86025 | 0.00015 |
| Phenylalanine and tyrosine catabolism | 6 | 8.95297 | 8.94E-05 | 4 | 9.31097 | 0.0086 |
| Phototransduction, and Guanylyl cyclase-activating protein 1 | 8 | 7.26661 | 0.0026 | 7 | 9.24331 | 0.00019 |
| Processing and activation of SUMO, and Sumo domain | 6 | -6.78139 | 0.0035 | 7 | -8.90086 | 0.0022 |
| Proteasome | 41 | -2.2322 | 0.0012 | 43 | -4.60545 | 6.82E-05 |
| Proteasome, and Poly [ADP-ribose] polymerase tankyrase-1 | 54 | -1.60821 | 0.0095 | 56 | -3.86209 | 0.00023 |
| Protein processing in endoplasmic reticulum, and Protein export | 60 | -1.91137 | 0.00065 | 50 | -5.84483 | 3.69E-06 |
| Pyruvate metabolism and Citric Acid (TCA) cycle, and 2-Oxocarboxylic acid metabolism | 55 | 2.98389 | 0.0029 | 57 | 5.12621 | 3.06E-05 |
| Recruitment of NuMA to mitotic centrosomes | 15 | -4.24008 | 0.0096 | 18 | -6.15582 | 0.0036 |

| | | | | | | |
|---|---|---|---|---|---|---|
| Recruitment of NuMA to mitotic centrosomes, and de novo centriole assembly involved in multi-ciliated epithelial cell differentiation | 25 | -4.32989 | 0.0011 | 21 | -6.55317 | 0.0039 |
| Regulation of Glucokinase by Glucokinase Regulatory Protein | 18 | -4.86645 | 0.00025 | 16 | -7.98739 | 5.67E-05 |
| Regulation of Insulin-like Growth Factor (IGF) transport and uptake by Insulin-like Growth Factor Binding Proteins (IGFBPs), and Platelet degranulation | 88 | 3.40951 | 4.51E-06 | 76 | 5.12088 | 3.22E-06 |
| Resolution of Sister Chromatid Cohesion, and chromosome segregation | 57 | -5.6423 | 4.30E-13 | 45 | -7.11549 | 8.00E-11 |
| Retinol metabolism, and The canonical retinoid cycle in rods (twilight vision) | 33 | 4.47411 | 0.00031 | 33 | 5.33601 | 0.00015 |
| RMTs methylate histone arginines | 12 | -5.04564 | 0.0031 | 11 | -7.84504 | 0.0021 |
| RMTs methylate histone arginines, and Requiem/DPF N-terminal domain | 23 | -4.16053 | 0.00054 | 21 | -5.60783 | 4.94E-05 |
| RNA Pol II CTD phosphorylation and interaction with CE | 11 | -5.82629 | 0.00063 | 11 | -8.48797 | 3.00E-04 |
| RNA Polymerase II Pre-transcription Events, and RNA polymerase II transcribes snRNA genes | 66 | -4.10059 | 1.32E-10 | 62 | -7.3494 | 3.88E-11 |
| RNA polymerase II transcribes snRNA genes | 18 | -4.32503 | 0.002 | 15 | -7.52169 | 0.00044 |
| RNA polymerase II transcribes snRNA genes, and Basal transcription factors | 46 | -4.69152 | 2.17E-10 | 43 | -7.95386 | 2.20E-10 |
| RNA polymerase II transcribes snRNA genes, and RNA Polymerase III Transcription Initiation From Type 1 Promoter | 86 | -3.58253 | 6.98E-10 | 79 | -6.19029 | 1.42E-09 |
| RNA Polymerase II Transcription Initiation | 28 | -4.92713 | 1.01E-06 | 28 | -8.18538 | 2.69E-08 |
| snRNP Assembly | 26 | -4.23475 | 0.00054 | 23 | -6.86906 | 0.0013 |
| snRNP Assembly, and Nuclear pore complex | 30 | -3.89415 | 0.0012 | 26 | -5.62157 | 0.0067 |
| snRNP Sm proteins, and PHF5-like | 18 | -6.45028 | 1.30E-06 | 15 | -8.63538 | 1.78E-06 |
| snRNP Sm proteins, and Spliceosome | 33 | -5.47253 | 2.96E-09 | 31 | -8.44539 | 1.16E-09 |
| Spliceosome | 58 | -5.00126 | 6.91E-14 | 53 | -7.74541 | 8.77E-12 |
| Telomere C-strand (Lagging Strand) Synthesis | 10 | -6.44312 | 0.00017 | 12 | -8.72293 | 3.06E-05 |
| Tryptophan/Indoleamine 2,3-dioxygenase-like, and Tryptophan 5-monooxygenase | 5 | 8.09881 | 0.0055 | 5 | 9.4682 | 0.00088 |

| | | | | | | |
|---|---|---|---|---|---|---|
| Ubiquitin-dependent degradation of Cyclin D1 | 27 | -3.41592 | 0.00014 | 30 | -6.05635 | 3.26E-05 |
| Voltage gated Potassium channels, and Phase 1 - inactivation of fast Na+ channels | 11 | 6.96504 | 6.00E-04 | 8 | 8.83231 | 0.00065 |
| Wnt signaling pathway, and Dickkopf, N-terminal cysteine-rich | 53 | -3.33858 | 0.00031 | 63 | -6.34922 | 5.50E-10 |
| Wnt signaling pathway, and Wnt signaling pathway | 72 | -2.97689 | 0.00017 | 76 | -6.22423 | 6.50E-11 |
